# Targeting the COP9 signalosome overcomes platinum resistance in ovarian cancer through two distinct genome stability mechanisms

**DOI:** 10.1101/2025.08.04.668450

**Authors:** Elena Lomonosova, Megan Loeb, Kevin Rodriguez, Lilian N van Biljon, Joshua Brill, Angela Schab, Margaret Minett, Jaeden Barron, Maya Bittner, Jiyoung Park, Rebecca Drexler, Carmen Sandoval, Alyssa Oplt, Eden Gallup, Prasanth K. Thuthika, Negar Sadeghipour, Sharon Wu, Matthew J Oberley, Brooke Sanders, Lindsay Kuroki, Carolyn McCourt, Andrea R. Hagemann, Premal Thaker, David Mutch, Matthew Powell, Ian S. Hagemann, Ma. Xenia G. Ilagan, Katherine Fuh, Priyanka Verma, Orlando D. Schärer, Nima Mosammaparast, Dineo Khabele, Mary M. Mullen

## Abstract

Tubo-ovarian high-grade serous carcinoma (HGSC) is a leading cause of gynecologic cancer mortality, largely due to the emergence of platinum resistance, which serves as the mainstay of chemotherapy. Here, we identify COPS5 as a therapeutic target and use an available small molecule inhibitor to overcome platinum resistance. A genetic screen for platinum-induced DNA damage in a platinum resistant ovarian cancer model identified COPS5 and COPS6, two components of the COP9 signalosome. Consistently, high COPS5 expression correlated with poor clinical outcomes in patients with HGSC. In both *in vitro* and *in vivo* experiments, COPS5 depletion sensitized ovarian cancer cells to carboplatin. A small molecule COPS5 inhibitor, CSN5i-3, synergized with carboplatin in homologous recombination-deficient and -proficient cells. This combination was also effective in xenografts and in a syngeneic mouse model of carboplatin-resistant HGSC. Importantly, we demonstrate that CSN5i-3 is selective for cancer cells, with patient-derived HGSC cells exhibiting up to 50-fold greater sensitivity to CSN5i-3 than benign cells. Finally, we show that genetic or small molecule inhibition of COPS5 impaired both nucleotide excision repair (NER) and interstrand crosslink (ICL) repair, leading to increased DNA platinum adducts. Mechanistically, this was due to increased ubiquitination and degradation of DNA-specific DNA binding protein 1 (DDB1) and other key NER and ICL repair proteins, consistent with the role of COPS5 in the regulation of these factors. Our findings highlight the importance of NER and ICL regulation in chemotherapy response and indicate that targeting COPS5 can enhance the efficacy of platinum-based chemotherapy in HGSC.

**One Sentence Summary:** COPS5 depletion or inhibition using a small molecule COPS5 inhibitor CSN5i-3 sensitizes high-grade serous carcinoma to platinum chemotherapy through downregulation of nucleotide excision repair and interstrand crosslink repair.

## INTRODUCTION

Over 85% of patients with tubo-ovarian high-grade serous carcinoma (HGSC) ultimately develop resistance to standard platinum chemotherapy (1–4). Platinum resistance is also a major therapeutic challenge in lung, bladder, head, and neck, colorectal, and triple-negative breast cancers. Platinum compounds act by forming covalent DNA adducts, leading to replication stress and DNA damage (5), and a primary contributor to resistance is upregulated activity of the DNA damage response (DDR) (6, 7). Among the DNA damage repair pathways, homologous recombination (HR) has received significant attention for its role in counteracting replication-induced breaks in response to platinum-induced lesions. Loss of HR capacity, as seen in BRCA1/2-mutated cancers, is associated with enhanced platinum sensitivity, while restoration or compensation of HR function contributes to resistance. However, most platinum-induced DNA lesions are initially repaired through the nucleotide excision repair (NER) or interstrand crosslink (ICL) repair pathways which remove bulky intrastrand and ICLs prior to replication (8, 9). Despite the central role of these pathways in lesion recognition and repair, NER and ICL repair have been relatively underexplored as therapeutic targets in platinum-resistant cancers. Growing evidence suggests that dysregulation or hyperactivation of NER or ICL repair components may critically influence platinum response and represents an untapped opportunity for therapeutic intervention (8). To improve outcomes for patients, strategies that overcome DDR-mediated resistance are urgently needed (10, 11).

Ubiquitin signaling is a key component of the DDR. It controls the recruitment and activity of repair proteins at sites of DNA lesions. Specifically, E3 ubiquitin ligases modify histones and repair factors to promote assembly of DDR complexes, while deubiquitinases (DUBs) remove ubiquitin to fine-tune repair timing and resolution. These interactions orchestrate DNA damage repair pathway choice, checkpoint activation, and genomic stability (12–15). Accordingly, interference of ubiquitin signaling can impact DNA repair, contributing to genomic instability and therapy response.

Therefore, to identify novel therapeutic targets, we focused on DUBs (12–15). In addition to their established role in DNA damage repair, deubiquitinating enzymes are attractive targets because of their structurally distinct catalytic domains, which can be selectively inhibited by small molecules. We performed a high-throughput siRNA screen targeting 98 deubiquitinating enzymes and identified two subunits of the COP9 signalosome complex as top candidates. The COP9 signalosome is a conserved complex comprised of eight subunits (COPS1-8) that removes neuronal precursor cell-expressed developmentally downregulated protein 8 (NEDD8) from cullin-RING E3 ubiquitin ligases (CRLs), which are required for E3 ligase activation (16, 17). Unlike most other DUBs, the COP9 signalosome catalytic activity is mediated by a metal-dependent protease domain within its MPN (JAMM) motif. This unique mechanism ensures specificity for NEDDylated CRLs (18, 19).

Here, we present evidence that loss of either of the two COP9 signalosome subunits, COPS5 or COPS6, leads to increased carboplatin-induced DNA damage in HGSC cell lines. Given the availability of a COPS5-specific inhibitor, we focused our work here on COPS5. Using patient-derived cells, patient-derived organoids, animal models, and clinical samples we systematically investigated the effect of COPS5 inhibition on platinum-adduct repair, and key genome stability mechanisms necessary to repair these lesions. These results provide the foundation for further investigation of deNEDDylation inhibition as a strategy to enhance platinum efficacy in HGSC.

## RESULTS

### Functional genetic screen reveals that depletion of COPS5 or COPS6 enhances platinum-induced DNA damage

Due to the role of ubiquitination in DNA damage response, we used short-interfering RNAs (siRNAs) to individually deplete nearly all (98) DUBs in the ovarian cancer cell line ES2. The cells were then treated with 1 µM cisplatin for 24 hours, and immunofluorescence was used to evaluate γH2AX, a marker of DNA damage (20–22) (*Fig. 1A*). Cisplatin was chosen over carboplatin for its higher potency and more consistent induction of DNA damage *in vitro* (23). We identified 17 candidates that, when depleted, increased the amount of cisplatin-induced γH2AX by at least three median absolute deviations above the median, indicating enhanced DNA damage. (*Fig. 1B, Supplementary Table S1*). Several of the identified candidates (USP7, OTUB1, OTUB2) have been reported to regulate DNA damage repair pathways (24), confirming the utility of our approach.

**Fig. 1.**
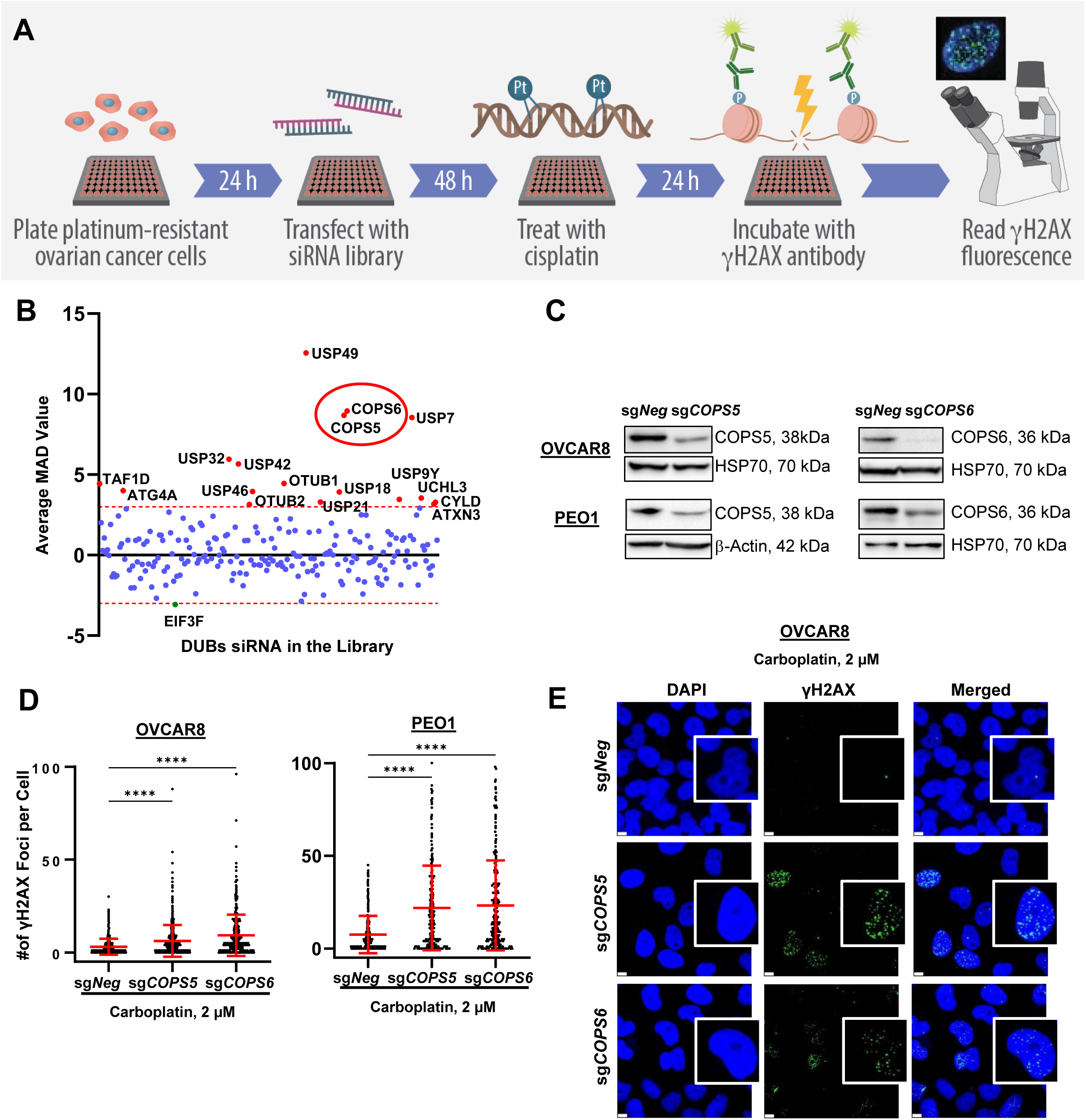
siRNA screen identifies COPS5 and COPS6 as mediators of platinum-induced DNA damage. **(A)** Flowchart of siRNA screen design. ES2 cells plated in 96-well plates were transfected with siRNA. Each gene was targeted by SMARTPools of four different siRNA targeting the same gene. Each plate also contained two wells of siCON (negative control), two wells of siDeath (positive control), and two wells of 10 µM cisplatin (positive control). Transfected cells were treated with vehicle or 1 μM cisplatin. Cells were stained with antibody to γH2AX to measure DNA damage. Images were collected on a laser-scanning fluorimeter, allowing quantification of γH2AX intensity. (**B**) Scatter plot of the average mean absolute deviation (MAD) (deviation in S.D. units from the plate median) for the screen. The dashed red lines denote MAD score=median±3 MAD. Candidates are defined as siRNAs with an average MAD difference from the relevant control of ≥ +3 or ≤ −3. Selected candidates are indicated by red dots. (**C**) COPS5 and COPS6 protein expressions in OVCAR8 and PE01 cells, as determined by Western blotting, with HSP70 as the loading control. (**D**) Quantification of γH2AX foci in sg*Neg,* sg*COPS5,* sg*COPS6* OVCAR8 and PE01 cells treated with carboplatin. (**E**) Representative images of γH2AX foci in sg*Neg,* sg*COPS5,* sg*COPS6* OVCAR8 cells treated with carboplatin. Scale bars, 5 µm. ****P <0.0001.

Two of the identified candidates were COPS5 and COPS6, which are known to form a heterodimer (25). These were prioritized based on their nuclear localization, high expression in HGSC, prior evidence of possible involvement in DDR, and the availability of a small-molecule inhibitor, which enhances their translational potential. We used the clustered regularly interspaced short palindromic repeats (CRISPR)/Cas9 system to individually knock out COPS5 and COPS6 in HGSC cell lines OVCAR8 and PE01, which was confirmed by immunoblotting (*Fig. 1C*) (26). To further explore their role in cell survival, we analyzed data from the Cancer Dependency Map, which reports genetic dependencies in cancer cell lines based on data from genome-wide CRISPR-Cas9 knockout screens. In HGSC cell lines, the Cancer Dependency Map scores for COPS5 and COPS6 were both below –1 (*Fig. S1A*), indicating that cells rely on those genes for survival. We then treated OVCAR8 and PE01 cells with carboplatin, the clinically used platinum drug, and found that it induced significantly more γH2AX foci in sg*COPS5* and sg*COPS6* cells than in controls (*Fig. 1D, E*). Together, these data suggest that COPS5 and COPS6 contribute to cancer cell viability and that their loss leads to increased platinum-induced DNA damage. Given that COPS5 and COPS6 are both subunits of the COP9 complex and that a COPS5 inhibitor is available (27), we focused the remainder of our experiments on COPS5.

### High COPS5 expression is associated with platinum resistance and poor prognosis in patients with HGSC

To directly evaluate the association between COPS5 and platinum chemotherapy resistance, we used eight patient-derived ovarian cancer cell lines (POVs) from patients with advanced-stage ovarian cancer. Western blots revealed that COPS5 expression significantly correlated with *in vitro* carboplatin resistance (Pearson r=0.81, *P*=0.02) (*Fig. 2A, Supplementary Fig. S1B*). Similarly, analysis of the Cancer Cell Line Encyclopedia dataset revealed that, in established cancer cell lines, resistance to cisplatin correlated with COPS5 protein expression (*Fig. 2B*). This suggested that COPS5 upregulation correlates with platinum resistance.

**Fig. 2.**
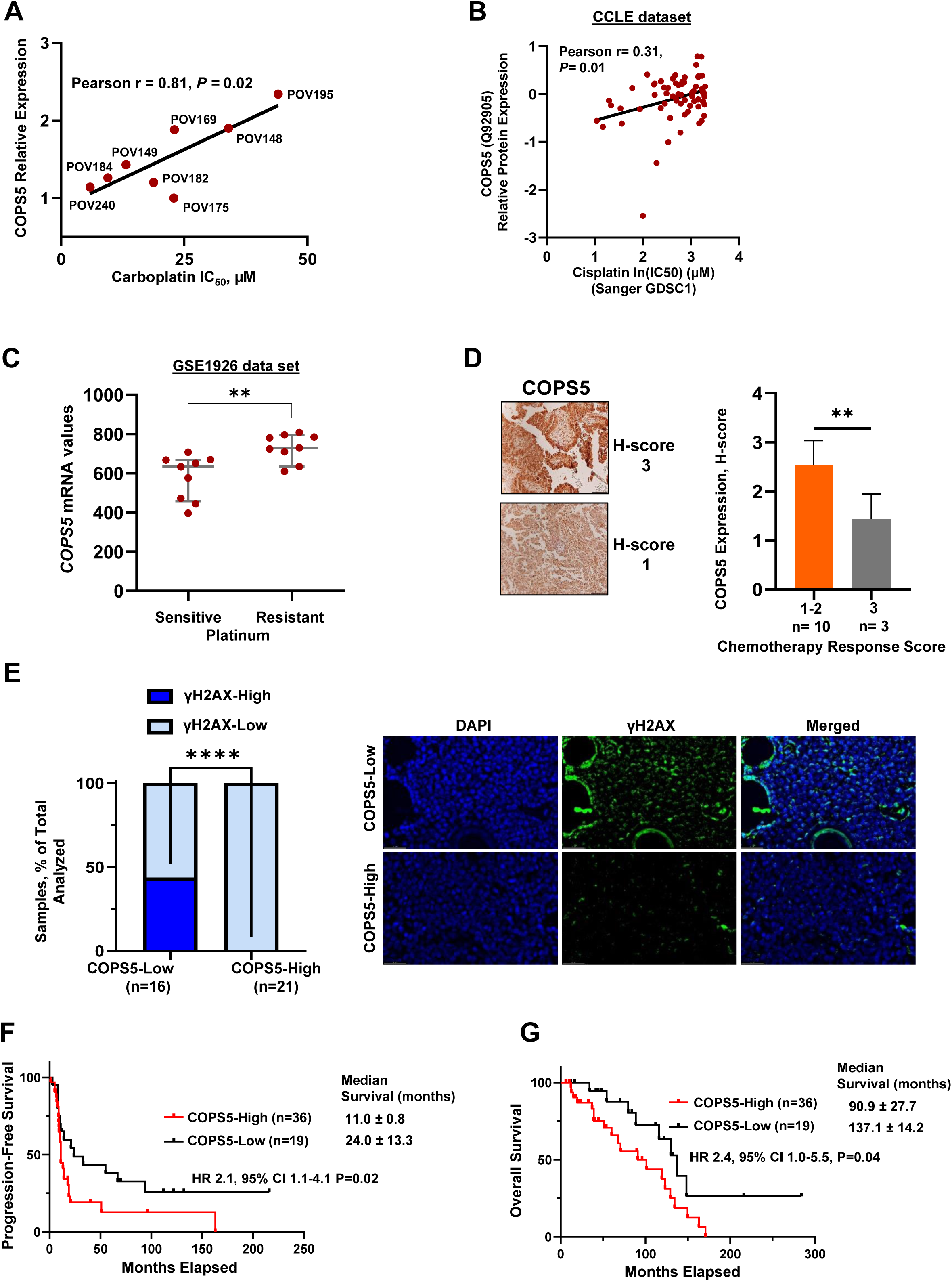
High COPS5 expression in patient-derived HGSC is associated with platinum resistance and poor prognosis. (**A**) Correlation of COPS5 protein expression and carboplatin IC50 values in POVs. (**B**) Correlation analysis of publicly available data of COPS5 protein expression and cisplatin IC_50_ values in cancer cell lines in the Cancer Cell Line Encyclopedia (CCLE) dataset. (**C**) Relative intensity of *COPS5* mRNA in HGSC cases described as platinum sensitive (n = 3 tumors examined in triplicate) or resistant (n = 3 tumors examined in triplicate). Data was analyzed by unpaired Student’s t test. (**D**) Representative immunohistochemistry images of HGSC samples stained for COPS5 (left). Scale bars, 50 μm. Bar graph depicting COPS5 expression in patients with partial (chemotherapy response score 1-2) and near complete response (chemotherapy response score 3) to platinum therapy (right). (**E**) Bar graph depicting the proportions of patients with high/low γH2AX and the expression of COPS5 (n = 37) (left) and representative images (right). Kaplan-Meier survival analysis comparing the (**F**) progression-free survival and (**G**) overall survival in patients with HGSC stratified by high vs. low COPS5 expression (n=55). Data were analyzed by unpaired Student’s t test [B, C, and E] and log-rank test [F and G].**P<0.01, ****P <0.0001.

To assess COPS5 in clinically relevant specimens, we took four approaches. First, we analyzed the publicly available Gene Expression Ontology dataset GSE1926 and found that platinum-resistant HGSCs (n= 9) expressed more COPS5 mRNA than did platinum-sensitive HGSCs (n=9) (*Fig. 2C*). Second, we analyzed COPS5 protein expression by immunohistochemistry in frozen fixed paraffin-embedded (FFPE) biopsy tissues from patients with stage III-IV HGSC. HGSC samples had been collected before and after three cycles of neoadjuvant chemotherapy and were assigned a chemotherapy response score (1 = poor response, 3 = nearly pathologic complete response) (28–30). HGSCs with a chemotherapy response score of 3 (n= 3) had significantly lower COPS5 expression than HGSCs with a chemotherapy response score of 1 or 2 (n=10) (*Fig. 2D*). We categorized these samples as COPS5-High (H-score ≥ 2) or COPS5-Low (H-score < 2) and evaluated the extent of DNA damage as indicated by the presence of γH2AX foci. HGSCs were defined as γH2AX-High if over 75% of cells had two or more γH2AX foci (31)). Whereas more than 40% of the COPS5-Low HGSCs were γH2AX-High, none of the COPS5-High HGSCs were γH2AX-High (*Fig. 2E*). Third, we measured COPS5 protein expression in HGSC samples from 55 patients in our Gynecologic Oncology Tissue Bank. 2 patients had FIGO stage II, 44 patients – stage III, 9 patients – stage IV, with median follow-up 54.2 +/− 58.5 months (6.0-284.00). Patients with COPS5-Low HGSCs had significantly longer progression-free *(Fig. 2F*) and overall (*Fig. 2G*) survival than patients with COPS5-High HGSCs. Finally, in 6,026 HGSC patient samples (Caris Life Sciences), patients whose samples had high COPS5 mRNA expression had significantly shorter overall survival than those with low COPS5 (39.6 vs. 41.5 months, HR 1.06, 95% CI 1.004-1.13, P=0.045) (*Supplementary Fig. S1C*). This difference was more pronounced in patients with *BRCA*-mutated HGSCs (47.9 vs. 58.8 months, HR 1.421, 95% CI 1.01-1.44, P=0.035) (*Supplementary Fig. S1D*). Together, these data indicate that high COPS5 expression corresponds to platinum resistance and poor survival outcomes in patients with HGSC.

### COPS5 depletion sensitizes ovarian cancer cells to carboplatin *in vitro* and *in vivo*

Given that COPS5 depletion increased platinum-induced DNA damage and its expression correlated with platinum resistance, we hypothesized that depleting COPS5 in ovarian cancer cells would enhance platinum sensitivity. To test this hypothesis, we treated sg*COPS5* and sg*Neg* OVCAR8 and PE01 cells with carboplatin and performed colony formation assays. The sg*COPS5* cells had significantly fewer colonies than sg*Neg* control cells, and this effect was reversed when we rescued COPS5 expression in sg*COPS5* PE01 cells by introducing a sgRNA-resistant COPS5 construct (*Fig. 3A, B*), indicating that the effect was specific to loss of COPS5. To further investigate whether COPS5 contributed to carboplatin response, we used siRNA to deplete COPS5 in several additional platinum-resistant ovarian cancer cell lines. siRNA-mediated depletion of COPS5 resulted in reduced cell viability from two to seven-fold upon carboplatin treatment (*Fig. 3C*).

**Fig. 3.**
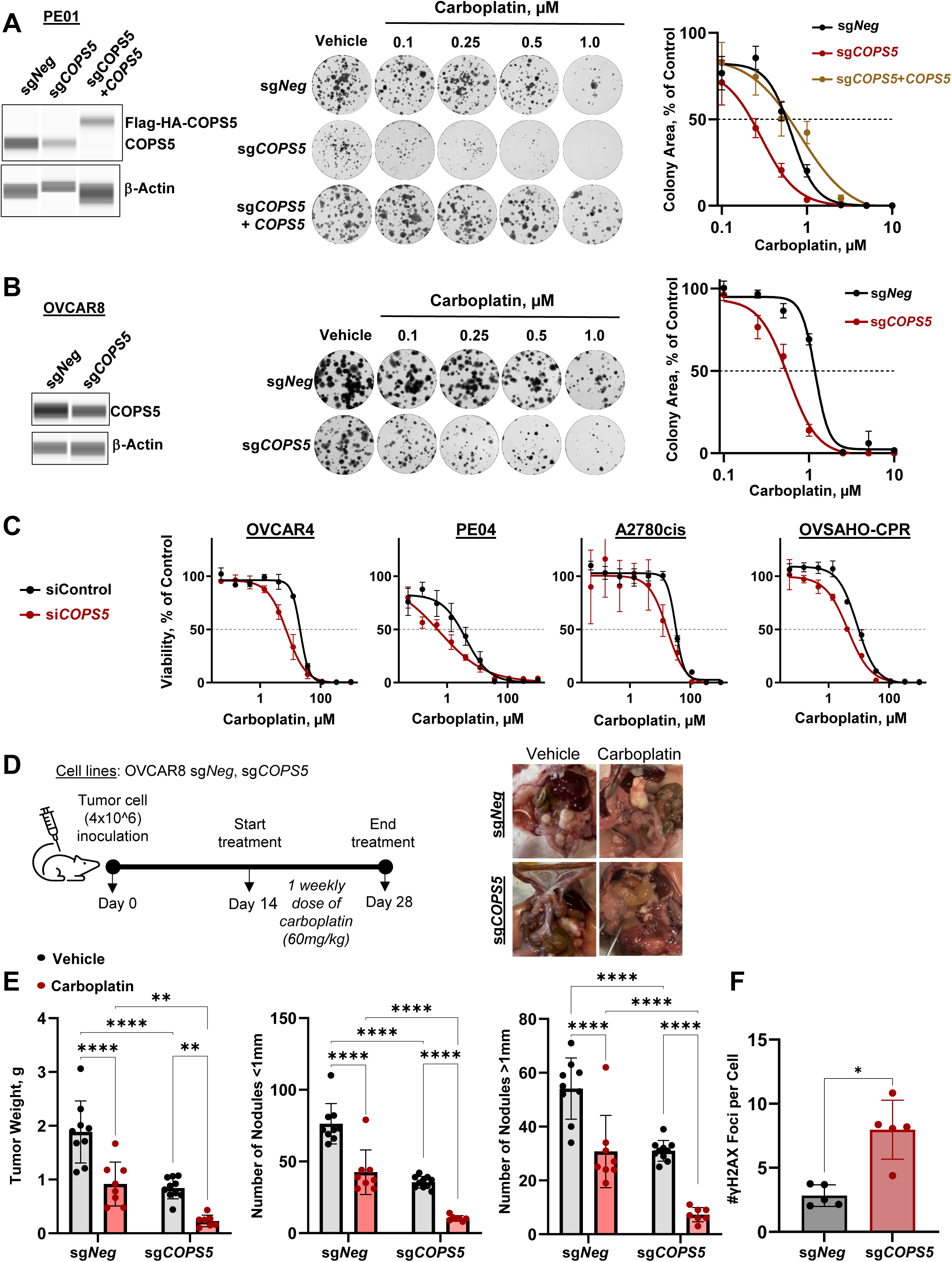
COPS5 depletion sensitizes HGSC to carboplatin *in vitro* and *in vivo*. Clonogenic survival for (**A**) sg*Neg*, sg*COPS5*, and sg*COPS5*+*COPS5* PE01 cells and (**B**) sg*Neg* and sg*COPS5* OVCAR8 cells treated with increasing doses of carboplatin. Representative images (left) and normalized survival (means ± SD, n = 3) fitted in log(inhibitor) vs. response Hill variable slope model (right). (**C**) Viability of indicated cells transfected with siControl and siCOPS5 and treated with carboplatin, as measured by MTS assay. (**D**) Schematic representation of the protocol for sg*Neg*, sg*COPS5* OVCAR8 xenograft model treated with single-agent intraperitoneal carboplatin and representative images. (**E**) Bar graphs showing tumor weight and number of nodules in each treatment group. (**F**) Bar graphs showing average number of γH2AX foci per cell in FFPE tumor samples of sg*Neg*, sg*COPS5* OVCAR8 xenografts (means ± SD, n = 3). Data was analyzed by Kruskal-Wallis test with Dunn’s correction for multiple comparisons. *P<0.05, **P<0.01, ****P <0.0001.

We next determined the effects of loss of COPS5 on tumor growth *in vivo*. OVCAR8 sg*Neg* and OVCAR8 sg*COPS5* cells were intraperitoneally implanted in nude mice. After 14 days, the mice were treated with vehicle or carboplatin (*Fig. 3D*). Tumor weight in sg*COPS5* xenografts was more than two times lower than in sg*Neg* xenografts at day 28 (0.84±0.2 vs 1.9±0.6 g, P <0.0001). Similarly, the number of tumor nodules was significantly lower in the sg*COPS5* group than in sg*Neg* (*Fig. 3D, E*). Importantly, sg*COPS5* tumors were significantly more sensitive to carboplatin than sg*Neg* counterparts as carboplatin treatment reduced sg*COPS5* tumor weight more than 3.5 times compared to up to 2-fold reduction in sg*Neg* tumors. Immunofluorescent analysis of FFPE tumor nodules in the vehicle-treated mice revealed significantly more γH2AX nuclear foci in the *sgCOPS5* group than in the sg*Neg* group, indicating higher endogenous DNA damage (*Fig. 3F*). Overall, these *in vivo* results were consistent with *in vitro* responses to carboplatin and confirmed that COPS5 depletion led to increased platinum sensitivity.

### The COPS5 inhibitor CSN5i-3 synergizes with carboplatin

We next evaluated a selective small-molecule inhibitor of COPS5, CSN5i-3 (27). CSN5i-3 binds to the JAB1/MPN/Mov34 metalloprotease domain of COPS5 and prevents it from deNEDDylating CRLs. Thus, CRLs remain in the active state, allowing them to ubiquitinate substrates to target them for proteasomal degradation. To confirm the reported CSN5i-3 specificity, we treated HGSC cells with varying concentrations of CSN5i-3 and observed accumulation of Cul4A in its NEDDylated, active state at nanomolar concentrations of CSN5i-3 (*Supplementary Fig. S2A*). Next, we confirmed that sg*COPS5* PE01 cells were less sensitive to CSN5i-3 than sg*Neg* PE01 cells (*Fig. 4A*). Similar results were found in the OVCAR8 cells (*Supplementary Fig. S2B).* To begin to assess whether CSN5i-3 would be toxic for non-malignant cells, we focused on fallopian tube cells, as HGSCs are presumed to originate in the fallopian tube epithelium (32). Using data from GSE10971, we found that *COPS5* mRNA expression was significantly higher in HGSCs than in fallopian tube epithelium cells (*Fig. 4B*). Furthermore, POVs were up to 50-fold more sensitive to CSN5i-3 than immortalized fallopian tube epithelium cells (FT282-hTERT) (*Fig. 4C*). Together, these observations confirmed the specificity of CSN5i-3.

**Fig. 4.**
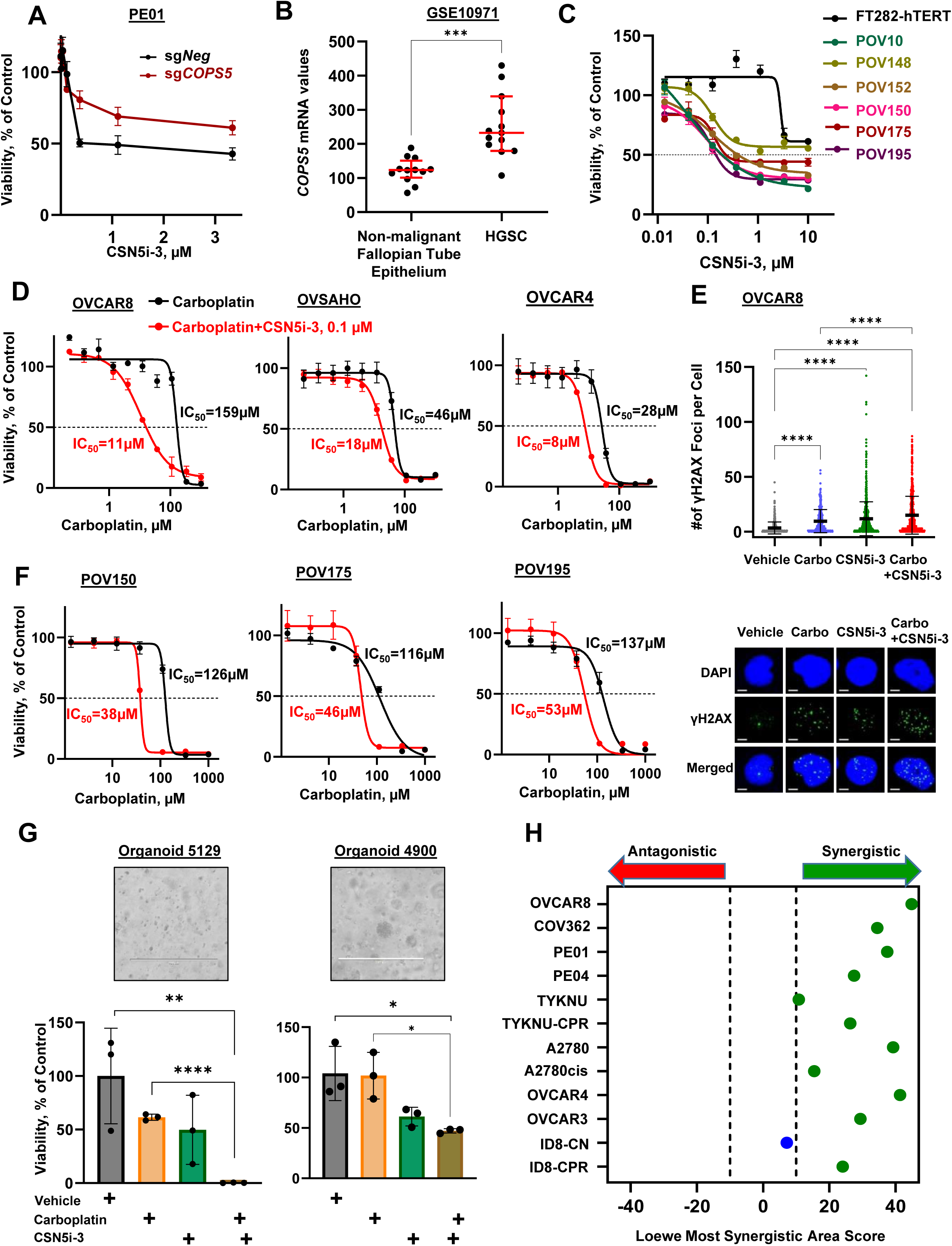
COPS5 inhibition with CSN5i-3 increases HGSC sensitivity to carboplatin. (**A**) Viability of sg*Neg* and sg*COPS5* PE01 cells treated with CSN5i-3, as measured by MTS assay (**B**) Relative intensity of *COPS5* mRNA in nonmalignant fallopian tube epithelium or HGSC, data generated from the Gene Expression OMnibus. Data were analyzed by unpaired Student’s t test. (**C**) Individual dose-response curves of CSN5i-3-treated indicated primary ovarian and immortalized fallopian tube cells, as measured by MTS assay. Dose–response curves represent normalized viability (means ± SD, n = 3) fitted in log(inhibitor) vs. response Hill variable slope model. Viability of indicated (**D**) established and (**F**) patient-derived ovarian cancer cells treated with increasing doses of carboplatin alone or in combination with 0.1 μM CSN5i-3, as measured by MTS assay. Dose–response curves represent normalized viability (means ± SD, n = 3) fitted in log(inhibitor) vs. response Hill variable slope model. IC_50_ values (means ± SD, n = 3) are indicated. (**E**) Quantification of γH2AX foci in OVCAR8 treated as indicated and representative images. Scale bars, 5 µm. (**G**) Results of viability assays of patient-derived organoids after treatment with CSN5i-3 plus carboplatin. Representative images and bar graphs depicting viability. Scale bars 100 μm. (**H**) Forest plot depicting Loewe most synergistic area score for carboplatin and CSN5i-3 combination in indicated cell lines. *P<0.05, **P<0.01, ***P<0.001, ****P <0.0001.

We hypothesized that CSN5i-3 would sensitize platinum-resistant ovarian cancer cells to carboplatin. To explore this, we performed viability assays in five HR-proficient, platinum-resistant ovarian cancer cell lines and found that CSN5i-3 enhanced carboplatin sensitivity by up to 15-fold (*Fig. 4D, Supplementary Fig. 2C).* As expected, CSN5i-3 increased platinum-induced DNA damage, evidenced by a significant increase in γH2AX foci in OVCAR8 cells from 9.64±10.5 average number of foci per cell in the carboplatin only group to 15.0±17.0 in the combination group, P<0.0001 (*Fig. 4E*). Consistent with our results in established cell lines, CSN5i-3 potentiated carboplatin toxicity in platinum-resistant HGSC POVs and patient-derived organoids (*Fig. 4F, G*). We next assessed drug synergy by using the Loewe additivity method across 12 HGSC cell lines (33), including HR-proficient (OVCAR8, COV362, PE04, TYKNU, A2780, OVCAR4, OVCAR3) and HR-deficient (PE01, ID8-CN) lines as well as models with *in vitro* (TYKNU-CPR, A2780cis) and *in vivo* (ID8-CPR) acquired platinum-resistance. CSN5i-3 demonstrated synergy with carboplatin in all lines except ID8-CN, which was highly platinum-sensitive at baseline (*Fig. 4H*). Together, these results demonstrate that CSN5i-3 enhances carboplatin efficacy in a variety of contexts including HR-proficient models.

### The addition of CSN5i-3 to carboplatin suppresses tumor growth and extends survival in a syngeneic mouse model of platinum-resistant HGSC

To test the utility of CSN5i-3 *in vivo* in immunocompetent mice, we generated a syngeneic model of platinum-resistant ovarian cancer. Although CSN5i-3 was originally developed as a human COPS5 inhibitor (27), the MPN+/JAMM motif of COPS5 is identical in *H. sapiens* and *M. musculus* (*Supplementary Fig. S3A*), and human and mouse COP9 subunits are 95-100% identical (Supplementary Table S2). Therefore, we reasoned that CSN5i-3 would also effectively target mouse COPS5. We generated novel chemo-resistant ID8-CPR model with *in vivo* acquired resistance to chemotherapy. We intraperitoneally injected ID8 cells into mice, treated the mice with carboplatin and paclitaxel, collected and plated ascites, and sorted the cells based on GFP expression so that only carcinoma cells remained in culture. We repeated this procedure over five passages, each with escalating doses of carboplatin and paclitaxel to generate carboplatin- and paclitaxel-resistant ID8 (ID8-CPR) cells (*Fig. 5A*). At the last passage, the ID8-CPR cells still expressed COPS5 and were three times more resistant to carboplatin than parental ID8 cells (*Supplementary Fig. S3B*). Next, we evaluated the combination of carboplatin and CSN5i-3 in this model (*Fig. 5B).* As expected, carboplatin monotherapy had only modest effects on ID8-CPR tumors (*Fig. 5C*). Conversely, the combination therapy significantly decreased intraperitoneal tumor growth, as measured by total tumor weight and number of tumor nodules (*Fig. 5C*). At the time of sacrifice, tumors from mice treated with CSN5i-3 expressed less COPS5 and Ki67 than mice from other groups (*Fig. 5D*). Finally, mice treated with carboplatin and CSN5i-3 had longer average overall survival (64.5 days) than mice treated with vehicle (14 days), carboplatin (26 days), or CSN5i-3 (14 days) (*P*<0.001) (*Fig. 5E*).

**Fig. 5.**
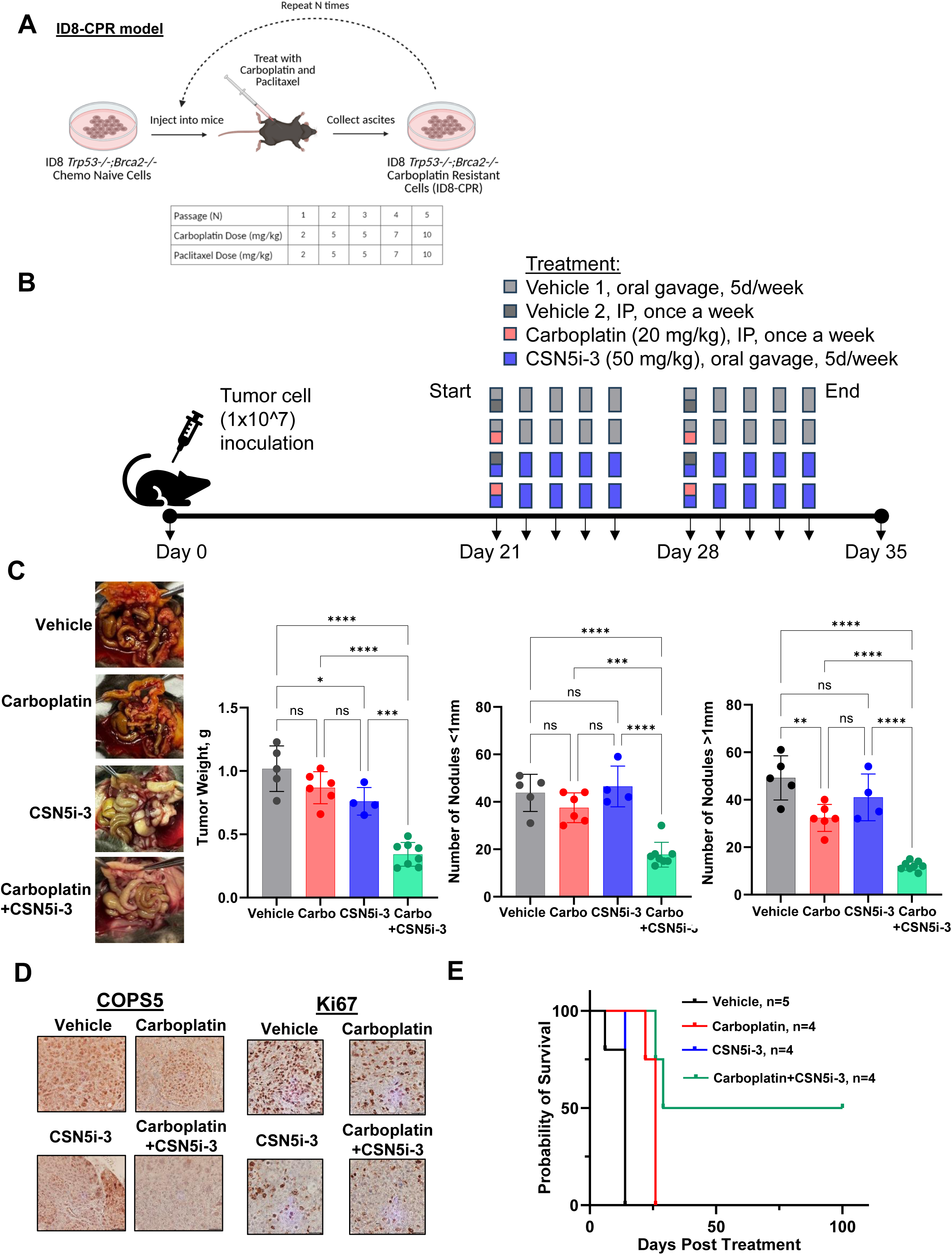
CSN5i-3 plus carboplatin reduces tumor growth and improves survival in a syngeneic mouse model of platinum-resistant HGSC. (**A**) A schematic illustration of generation of carboplatin and paclitaxel-resistant ID8 (ID8-CPR) cells (created with Biorender.com). (**B**) Schematic representation of the protocol for ID8-CPR syngeneic model and experimental treatment strategy. ID8-CPR cells were inoculated intraperitoneally (IP) into mice on day 0. Beginning on day 21, vehicles, carboplatin, and CSN5i-3 were injected at the indicated doses and schedules. (**C**) Representative images of tumors in ID8-CPR model. Bar graphs showing tumor weight and tumor nodules at the end of treatment (mean±SD). (**D**) Representative immunohistochemistry images showing COPS5 and Ki67 staining in treated tumors. Scale bar, 50 μm. (**E**) Kaplan–Meier analysis comparing overall survival of mice injected with ID8-CPR cells treated with vehicle, carboplatin, CSN5i-3, or carboplatin plus CSN5i-3. ***P<0.001, ****P <0.0001.

### COPS5 depletion downregulates NER

Given that CSNi-3 enhanced platinum sensitivity in an HR-deficient setting, it suggests that the impact of COPS5 is beyond homology directed repair of DNA double strand breaks. To begin exploring the mechanism by which COPS5 inhibition sensitizes ovarian cancer cells to carboplatin, we first evaluated transcriptomic data from ovarian cancer samples (Caris Life Sciences dataset). Although COPS5 is primarily known to regulate protein stability through post-translational mechanisms, this initial mRNA analysis was used to identify DNA repair pathways associated with COPS5 expression in patient tumors. We compared cancers with high (> 75%) versus low (<25%) COPS5 mRNA expression and found that those with high expression showed significant enrichment of DNA repair pathways (*Supplementary Fig. S4A*). Gene set enrichment analysis (GSEA) revealed that this upregulation was primarily driven by increased expression of genes involved in nucleotide excision repair (NER), the Fanconi anemia (FA) pathway, and homologous recombination (HR) (*Supplementary Fig. S5A, S5B*). This finding was intriguing because platinum-DNA adducts lead to intrastrand crosslinks, which are primarily repaired by NER (34, 35), and ICLs, which are processed via FA and HR (35, 36). NER consists of two sub pathways: global genome NER (GG-NER), which removes DNA damage from the entire genome, and transcription-coupled NER (TC-NER), which corrects lesions located on actively transcribed genes (37, 38). In previous work, depletion of COPS5 in fibroblasts and embryonic stem cells led to defects in both GG-NER and TC-NER (26, 39).

To determine whether loss of COPS5 leads to reduced NER in HGSC, we treated *sgCOPS5* OVCAR8 and PE01 cells with UVC irradiation. UVC irradiation is known to induce DNA damage that is primarily repaired by NER. Following UVC irradiation, sg*COPS5* OVCAR8 and PE01 cells formed significantly fewer colonies than sg*Neg* cells (*Fig. 6A*). Similarly, CSN5i-3 treatment sensitized OVCAR8 and PE01 cells to UVC irradiation (*Fig. 6B*). Next, we treated cells with high-dose formaldehyde, which causes DNA-protein crosslinks (40) that are resolved by a pathway involving the TC-NER proteins CSA and CSB (41–43). We found that sg*COPS5* OVCAR8 and PE01 cells were more sensitive to formaldehyde than were sg*Neg* cells (*Fig 6C*). Likewise, CSN5i-3 treatment increased sensitivity to formaldehyde (*Fig. 6D*). These results suggest that COPS5 contributes to both TC- and GC-NER in ovarian cancer cells. Given that carboplatin induced lesions necessitate TC-NER and GC-NER, it is likely that these pathways underlie COPS5-mediated platinum resistance in HGSC.

**Fig. 6.**
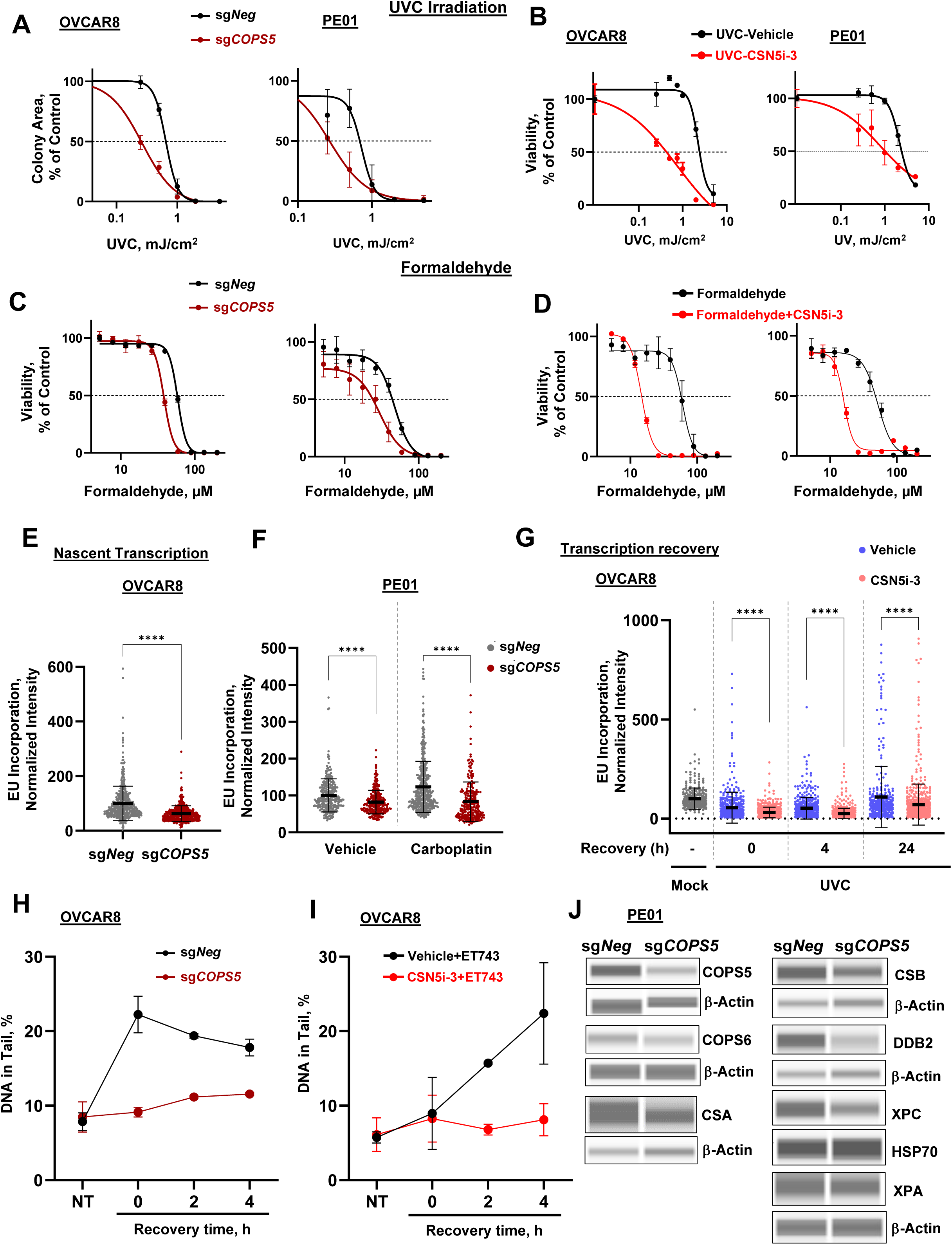
COPS5 inhibition sensitizes ovarian cancer cells to carboplatin via downregulation of NER. (**A**) Viability of indicated cells treated with increasing doses of UVC alone or (**B**) in combination with 0.1 μM CSN5i-3, as measured by MTS assay. Dose–response curves represent normalized viability (mean ± SD, n = 3) fitted in log(inhibitor) vs. response Hill variable slope model. (**C**) Viability of indicated cells treated with increasing doses of formaldehyde alone or (**D**) in combination with 0.1 μM CSN5i-3, as measured by MTS assay. Dose–response curves represent normalized viability (mean ± SD, n = 3) fitted in log(inhibitor) vs. response Hill variable slope model. (**E**) Quantification of nascent transcripts labelled with ethynyl-uridine (EU) in OVCAR8 sg*Neg* and sg*COPS5* cells. (**F**) Scatter plot (left) and representative fluorescence images (right) showing nascent transcripts labelled with EU and assessed after 24 hours after vehicle or carboplatin administration in PE01 sg*Neg* and sg*COPS5* cells. Data points indicate normalized nuclear EU intensity per cell. Data are mean ± SD, n > 200 cells. Scale bar, 5 μm. DAPI, 4′,6-diamidino-2-phenylindole. (**G**) Scatter plot showing quantification of recovery of RNA synthesis in OVCAR8 cells treated with vehicle or 0.1 μM CSN5i-3 and then mock- or UV-irradiated (2 mJ/cm2). mRNA was labeled with EU at the indicated time points post-UV. EU signal was quantified by ImageJ and relative integrated densities, normalized to mock-treated level set to 100%, are reported on the graph, n > 200 cells per condition. Black bars indicate mean integrated density. Error bars ± SD. (**H**) Summary and statistical analysis of ssDNA breaks analyzed by alkaline COMET chip assay in OVCAR8 sg*Neg* and sg*COPS5* cells treated with trabectedin (50 nM, 2 h) and allowed to recover for up to 4 h. Mean ± SEM of at least three biological replicates. (**I**) Summary and statistical analysis of ssDNA breaks analyzed by alkaline COMET chip assay in OVCAR8 sg*Neg* and sg*COPS5* cells treated CSN5i-3 (2 h) and then with trabectedin (50 nM, 2 h) and allowed to recover for up to 4 h. Mean ± SEM of at least three biological replicates. Each dot represents DNA in tail (%) of a comet analyzed. (**J**) Jess Simple Western blots showing indicated proteins in PE01 sg*Neg* and sg*COPS5* cells with β-actin as the loading control. Statistical significance was calculated by two-tailed paired t-test (**E, G**) and one-way ANOVA (**F, I**) with Welch’s test, or ordinary two-way ANOVA with Dunnett’s multiple comparisons test. ****P < 0.001.

DNA damage caused by UVC irradiation and platinum-based chemotherapy (44–46) leads to transcriptional shutdown, requiring NER-mediated repair of transcription-blocking DNA lesions (37, 47, 48). We hypothesized that in COPS5-deficient cells, carboplatin-induced DNA damage that remains unrepaired by NER would lead to greater inhibition of global transcriptional activity. To test this hypothesis, we quantified nascent RNA synthesis by examining nuclear incorporation of the modified RNA precursor 5-ethynyluridine (EU) in sg*Neg* and sg*COPS5* cells. In both vehicle- and carboplatin-treated conditions, nascent RNA production was lower in sg*COPS5* OVCAR8 and PE01 cells than in sg*Neg* cells (*Fig. 6E, F*). Similarly, nascent RNA synthesis was downregulated in cells upon treatment with CSN5i-3 (*Supplementary fig. S6A*).

Recovery of RNA synthesis (RRS) after DNA damage induction is an indirect indicator of TC-NER activity (49). We therefore analyzed RRS after UVC irradiation in OVCAR8 and PE01 cells-treated with CSN5i-3 by pulse-labeling with EU at various time points after UVC irradiation. UVC irradiation inhibited transcription, which recovered fully after 24 hours in control cells (*Fig. 6G, Supplementary Fig. S6B*). In contrast, transcription recovery in CSN5i-3-treated cells did not return to baseline after 24 hours, indicating that COPS5 contributes to transcription recovery after UV irradiation.

We next examined the effects of COPS5 knockdown or inhibition on the DNA-damaging activity of trabectedin (Ecteinascidin 743, ET743). Trabectedin is an anticancer drug that is more toxic to cells with active TC-NER. When TC-NER acts on trabectedin-DNA adducts, it causes persistent single-strand DNA (ssDNA) breaks that underlie its DNA repair dependent toxicity (50, 51). Therefore, functional TC-NER can be evaluated by treating cells with trabectedin and performing alkaline CometChip assays to measure ssDNA break formation (51, 52). If TC-NER is disabled, there will be fewer ssDNA breaks. We found that after treatment with trabectedin, sg*Neg* cells had more ssDNA breaks than sg*COPS5* OVCAR8 cells (*Fig. 6H*). Similarly, trabectedin-induced ssDNA breaks in sg*Neg* OVCAR8 (*Fig. 6I*), U2OS, and RPE-1 cells (*Supplementary Fig. S6D*) were decreased following COPS5 inhibition with CSN5i-3. CSN5i-3 had no effect on trabectedin-induced ssDNA breaks in sg*COPS5* OVCAR8 cells, confirming its specificity (*Supplementary Fig. S6C*). Overall, these data support a key function of COPS5 in TC-NER.

Previous studies suggested that COPS5 depletion leads to decreased expression of a key mediator of HR, RAD51 (26, 53). However, RAD51 expression alone does not consistently reflect repair capacity (31, 54). To assess the impact of COPS5 loss on DNA repair dynamics, we measured RAD51 nuclear foci formation upon treatment of OVCAR8 cells with CSN5i-3 alone or in combination with carboplatin. We observed a compensatory increase in RAD51 foci after CSN5i-3 treatment alone and in combination with carboplatin (*Supplementary Fig. S6E*). These results suggest that the increased dsDNA breaks observed after carboplatin treatment in sg*COPS5* cells (*Fig. 1E*) cannot be explained by an inhibitory effect on RAD51 loading.

We hypothesized that COPS5 inhibition abrogated NER by affecting the abundance of NER proteins through ubiquitination and proteasomal degradation. We found that the abundance of key proteins in both TC-NER (CSA, CSB) and GG-NER (DDB2, XPC) were reduced in sg*COPS5* PE01 cells (*Fig. 6J*). Similar results were observed when OVCAR8 cells were treated with CSN5i-3 or CSN5i-3 plus carboplatin (*Supplementary Fig. S6F*). COPS5 and the COP9 signalosome are involved in control of the ubiquitin-mediated degradation of ∼20% of cellular proteins (17, 55). Additionally, COPS5, originally termed JAB1, functions as a transcriptional co-activator of c-Jun, a component of the activator protein-1 complex (56, 57). Given our observation that COPS5 depletion significantly reduced transcription (*Fig. 6*), we sought to confirm that protein abundance differences were driven by proteasomal degradation and not transcriptional differences. Thus, we treated OVCAR8 cells with CSN5i-3 plus the proteasome inhibitor MG-132 (58). MG-132 treatment resulted in partial rescue of key TC-NER and GG-NER proteins including DDB2 and CSA (*Supplementary Fig. S6G*). These data support a model in which CSN5i-3 disrupts ubiquitin-dependent degradation of repair proteins involved in NER.

### COPS5 inhibition downregulates ICL repair

While NER is essential for repairing platinum-induced intrastrand crosslinks, NER deficiency alone results in only mild cisplatin sensitivity. This is likely because intrastrand lesions are less genotoxic than ICLs (59). Given the marked increase in cisplatin sensitivity observed with COPS5 inhibition, we hypothesized that COPS5 may also play a role in additional repair pathways critical for ICL resolution. To begin to evaluate this, we performed tandem ubiquitin-binding entity (TUBE)-based enrichment followed by mass spectrometry using the LifeSensors platform (*Figure 7A*). Considering the transient nature of ubiquitinated proteins, we treated samples with CSN5i-3 in the presence of a low dose of MG132 to better capture ubiquitinated proteins. A total of 3205 ubiquitinated proteins were identified (*Supplementary Table S3).* GSEA (60) revealed prominent enrichment of ubiquitinated proteins in processes of cellular response to stress, protein catabolic process, cell cycle, post-translational modification, DDR, and DNA damage repair (*Supplementary Figure S7A, Supplementary Table S4).* We observed increased ubiquitination of all subunits of the COP9 signalosome, suggesting potential autoregulatory feedback on signalosome function or destabilization of the complex in response to COPS5 inhibition (*Supplementary Fig. S7B*). We identified increased ubiquitination of DDR genes (*Figure 7B, Supplementary Figure S7C).* Consistent with our previous findings that CSN5i-3 downregulates NER, we found increased ubiquitination of key NER factor DDB1. DDB1 is a core component of the CUL4– DDB1 E3 ubiquitin ligase complex, which regulates the stability and activity of multiple proteins involved in DNA damage recognition and repair. Downregulation of DDB1 expression upon CSN5i-3 treatment was confirmed by western blot (*Supplementary Fig. S6H*). We also found modulation of ubiquitination of several ICL repair factors, including proteins in the Fanconi Anemia pathway (*Figure 7B*). We demonstrated the reduced expression of FANCA, FANCI, and FANL in OVCAR8 cells treated with CSN5i-3 by western blot analysis (*Figure 7C*).

**Fig. 7.**
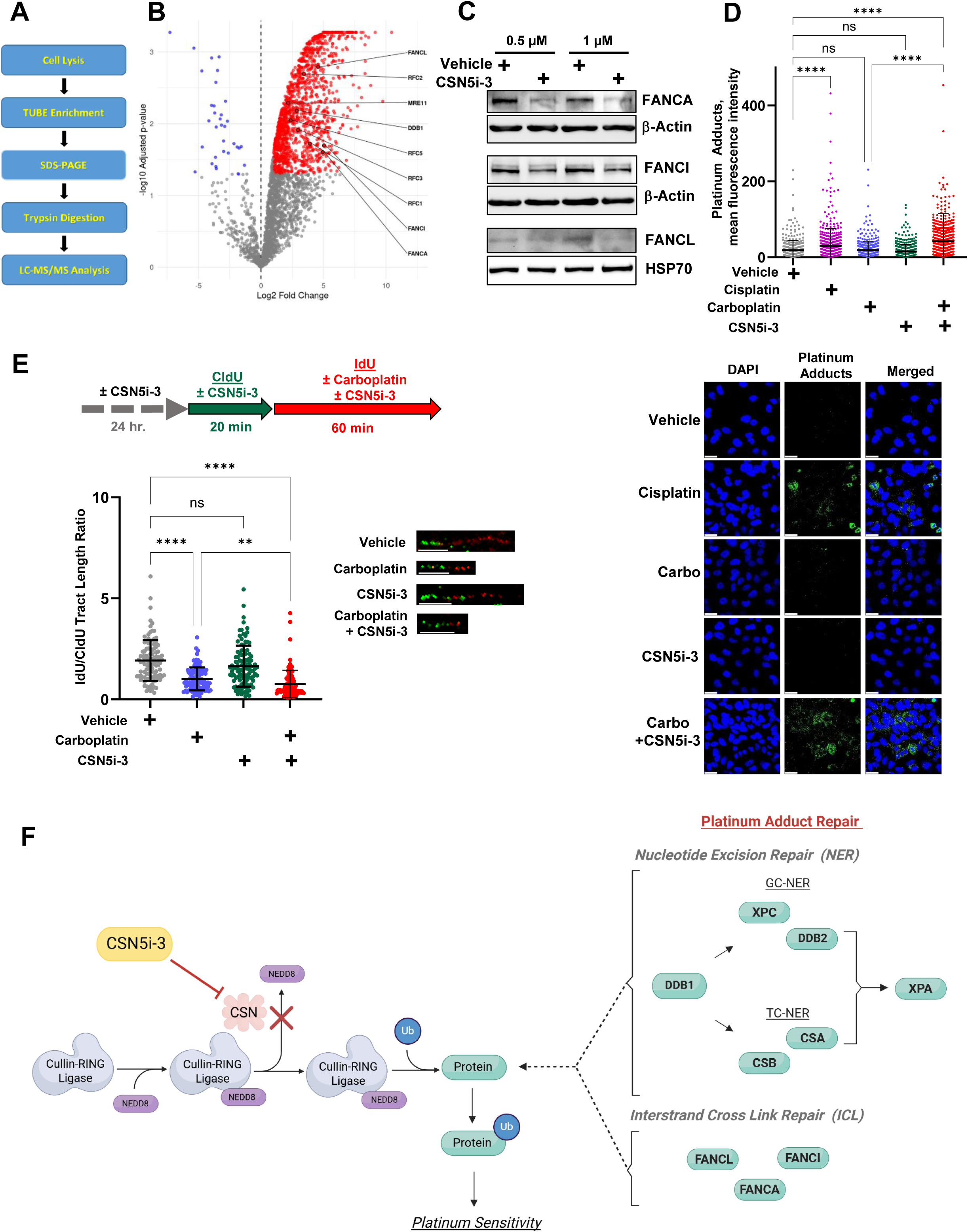
COPS5 inhibition reduces expression of proteins involved in NER. (**A**) Experimental flowchart for ubiquitinome profiling by mass spectrometry. Cells were treated with vehicle, CSN5i-3, or CSN5i-3 and MG132, followed by lysis and enrichment of ubiquitinated proteins using a tandem ubiquitin-binding entity (TUBE)-based pulldown. Enriched proteins were digested and subjected to liquid chromatography–tandem mass spectrometry (LC-MS/MS). Quantitative proteomics analysis was performed to identify differentially ubiquitinated proteins between treatment groups. (**B**) Volcano plot of differentially expressed ubiquitinated proteins following CSN5i-3+MG132 treatment. Proteins involved in DNA damage response pathway are highlighted. (**C**) FANCA, FANCI, FANCL protein expression in OVCAR8 treated with CSN5i-3, as determined by Western blotting, with β-actin as the loading control. (**D**) Quantification of platinum-DNA adducts after treatment with vehicle, cisplatin 10 µm, carboplatin 2µm, 0.1 μM CSN5i-3 for 24 hours, or CSN5i-3 and carboplatin. Error bars ±SD, n = 3 replicates. (**E**) DNA fiber assay analysis with schematic and representative images of the labeling procedure and quantitative analysis of IdU/CldU tract length ratio upon treatment with vehicle, 500 μM carboplatin, 0.1 μM CSN5i-3, or combination. CldU, 5-chloro-2’-deoxyuridine; IdU, 5-iodo, 2’-deoxyuridine; **P < 0.01, ***P < 0.001 by Kruskal-Wallis test. Scale bars are 5 μm. (**F**) Model depicting the proposed mechanism by which CSN5i-3 enhances platinum adduct repair via inhibition of the COP9 signalosome, leading to sustained neddylation of cullin-RING ligases leading to excessive degradation of key DNA repair proteins, thus disrupting NER and ICL repair pathways

To further substantiate the role of COPS5 in platinum-DNA adducts repair, we directly measured platinum– DNA adduct accumulation using an anti-cisplatin modified DNA antibody following treatment with CSN5i-3. Compared to carboplatin alone, cells treated with the combination of carboplatin and CSN5i-3 exhibited significantly increased levels of persistent platinum adducts as assessed by immunofluorescence for platinum adducts (*Fig. 7D*). Measuring replication fork progression in platinum-treated cells provides a functional readout of how effectively cells manage DNA crosslinks, including ICLs. Specifically, impaired ICL repair would be expected to lead to persistent platinum-DNA adducts and consequent replication fork stalling. To assess the impact of COPS5 inhibition on replication fork processivity and replication efficiency, we performed DNA fiber analysis, which showed reduction of replication fork speed after combination treatment with carboplatin and CSN5i-3 (*Figures 7E*). Collectively, these data support the conclusion that CSN5i-3 results in increased NEDDylation of CRLs resulting in increased ubiquitination and subsequent degradation of key NER and ICL repair factors. This impairs adduct resolution and enhances DNA damage accumulation, resulting in slowing of fork progression upon encountering DNA damage (*Figure 7F*).

Lastly, given that COPS6 functions as a heterodimeric partner of COPS5 within the COP9 signalosome, we investigated whether COPS6 similarly regulates platinum sensitivity. Depletion of COPS6 phenocopied the effects of COPS5 depletion, further supporting the role of the COP9 signalosome in modulating platinum chemotherapy response (Figure 8, Supplementary Fig. S4B, S7D-G).

**Fig. 8.**
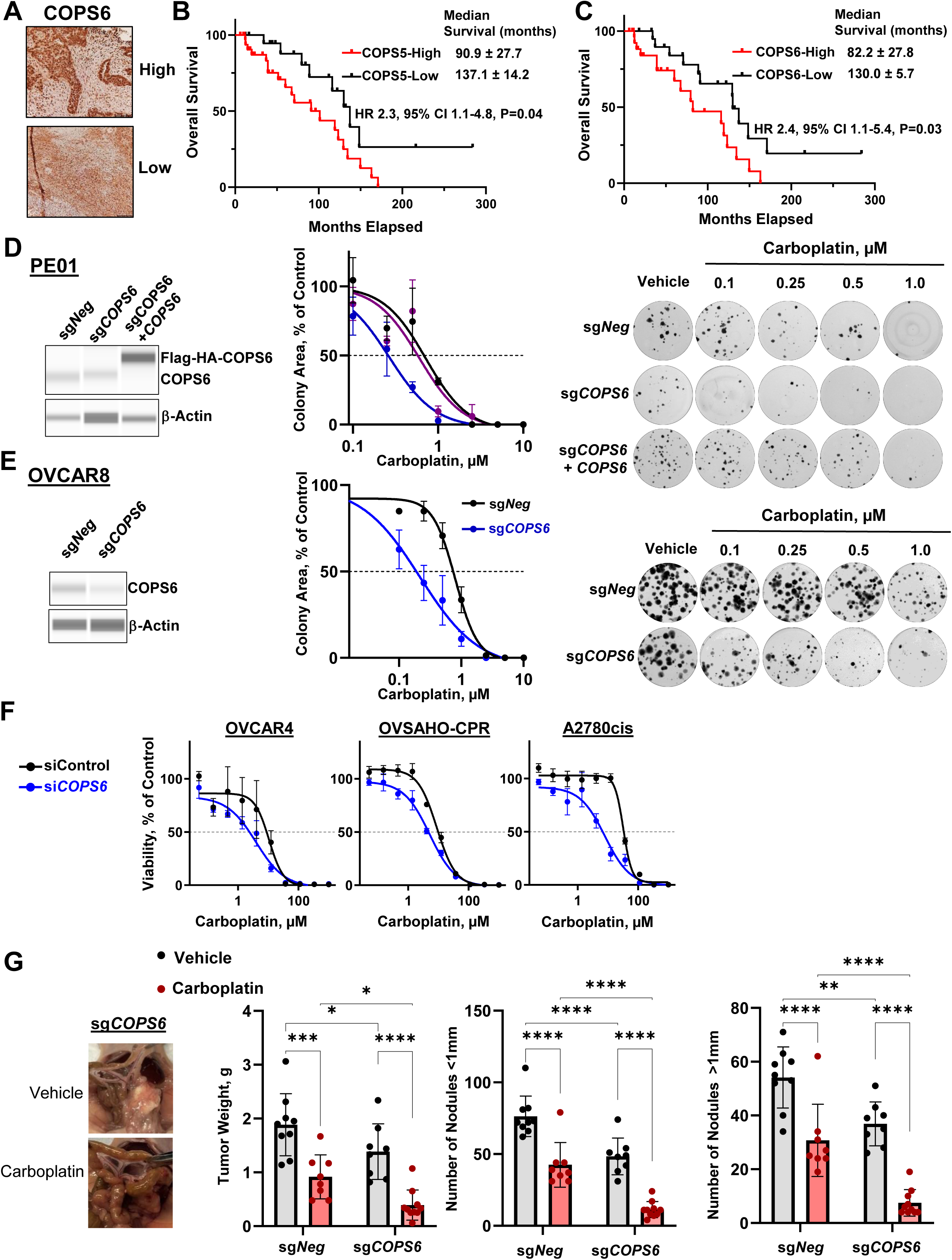
COPS6 mediates platinum sensitivity. **(A)** Representative immunohistochemistry images of HGSC samples stained for COPS6. Kaplan-Meier survival analysis comparing the (**B**) progression-free survival and (**C**) overall survival in patients with HGSC stratified by high vs. low COPS6 expression (n=55). **(D)** sg*Neg*, sg*COPS6*, and sg*COPS6*+*COPS6* PE01 cells and **(E)** sg*Neg* and sg*COPS6* OVCAR8 cells treated with increasing doses of carboplatin. Representative images (right) and normalized survival (means ± SD, n = 3) fitted in log(inhibitor) vs. response Hill variable slope model (left). (**F**) Viability of indicated cells transfected with siControl and siCOPS6 and treated with carboplatin, as measured by MTS assay. **(G)** Representative images of sg*Neg*, sg*COPS5* OVCAR8 xenograft model treated with single-agent intraperitoneal carboplatin (left), and bar graphs showing tumor weight and number of nodules in each treatment group (right). Data were analyzed by unpaired Student’s t test [A], log-rank test [B and C], and Kruskal-Wallis test with Dunn’s correction for multiple comparisons [D, E, F, and G]. *P<0.05, **P<0.01, ****P <0.0001.

## DISCUSSION

Platinum-based drugs are key in ovarian cancer treatment, but resistance and toxicity pose major challenges (61–66). Here, we show that genetic depletion of COPS5 and COPS6 increases platinum sensitivity in HGSC. High COPS5 expression was associated with increased platinum resistance and worse clinical outcomes in patients with ovarian cancer, consistent with its prognostic role in other cancers (25, 67, 68). We show that inhibition of COPS5 increases platinum sensitivity in multiple patient-derived models including HR-proficient, HR-deficient, and platinum-resistant cells as well as organoids. Additionally, we show that *in vivo* inhibition of COPS5 sensitizes a syngeneic mouse model of HGSC to platinum. Finally, we show that targeting COPS5 enhances the efficacy of platinum-based chemotherapy by promoting stability of proteins involved in NER and FA pathways. Notably, inhibition of COPS5 did not enhance carboplatin sensitivity of non-cancerous hTERT FT282 fallopian tube cell line. Together, this work provides a strong foundation for the clinical translation of COPS5 inhibition as a therapeutic strategy to overcome platinum-resistance in HGSC.

NER and ICL are the primary pathway for repairing cisplatin-induced DNA adducts (69), and expression of key NER proteins correlates with platinum response (70–73). However, pharmacologically targeting NER and/or ICL has proved challenging (74–77). Our study reveals that COPS5 depletion or pharmacological inhibition promotes proteasomal degradation of key NER and ICL proteins, reducing NER and ICL repair capacity and increasing platinum-induced cell death (*Figure 7F*).

COPS5 regulates NER protein stability through CRL-mediated degradation, and its inhibition disrupts this process. COPS5 normally removes NEDD8 modifications which prevents excessive CRL activation, auto-ubiquitination, and degradation of target proteins (17, 18, 78, 79). CRL4 complexes, such as CUL4^CSA^ and CUL4^DDB2^, regulate key NER factors, including DDB1, CSA, and DDB2 (39, 80), all of which were observed to be degraded upon COPS5 depletion despite typically being stabilized in response to DNA damage. This degradation is likely a direct consequence of deNEDDylation inhibition. Presumably substrates of these CRL4 complexes would be spared from degradation given the absence of CRL4 complexes. However, CSB and XPC were also degraded upon COPS5 depletion despite being substrates of CUL4^CSA^ and CUL4^DDB2^ (81, 82). This suggests that loss of COPS5 leads to hyperactivation and dysregulated ubiquitination of CRL substrates, bypassing canonical regulation and triggering aberrant protein turnover. Notably, XPC ubiquitination has been linked to protein stabilization rather than degradation, which may be disrupted by deNEDDylation inhibition (82, 83). It is also possible that differential turnover of CRL substrates in the absence of COP9 activity contributes to these effects, as the sensitivity of CRLs to deNEDDylation is highly context dependent. Given that CRLs regulate both GG-NER and TC-NER, COPS5 likely plays a central role in coordinating DNA repair protein turnover to preserve transcriptional fidelity.

We also found that COPS5 inhibition disrupts ICL repair which is a critical DDR pathway activated by platinum-induced DNA damage. Our data show that COPS5 inhibition increases ubiquitination and degradation of multiple FA proteins. Very little is known about NEDD8-dependent regulation of FA proteins stability (84), and to the best of our knowledge, this is the first observation that FA proteins are controlled by CSN complex. Further studies are needed to identify mechanistic aspects of this regulation. ICL repair involves the coordinated action of the FA pathway, NER, and HR. Destabilization of ICL repair by CSN5i-3 results in increased platinum adduct accumulation, replication stress, and increased DNA damage, thereby enhancing platinum sensitivity. Together, these findings suggest COPS5 is a central post-translational regulator of ICL repair fidelity and genome integrity.

Targeting COPS5 has strong potential as a therapeutic strategy for treating platinum-resistant ovarian cancer for several reasons. First, COPS5 inhibition sensitized ovarian cancer cells to carboplatin without significantly affecting normal fallopian tube epithelial cells, suggesting a favorable therapeutic index. Second, targeting the NEDDylation/deNEDDylation cycle has promise in cancer treatment (85). Inhibition of the NEDD8-activating enzyme with MLN4924 (Pevonedistat) slowed ovarian cancer proliferation in various preclinical models and is currently being evaluated in clinical trials (85–88). However, whereas MLN4924 blocks the entire NEDDylation process, leading to broad cytotoxic effects (89), COPS5 inhibition affects only NEDDylated CRLs rather than all NEDDylated proteins (27). Thus, a COPS5 inhibitor may have greater specificity and fewer side effects than MLN4924. Third, COPS5 is an attractive target given its structural and enzymatic properties. Deubiquitinases generally contain well-defined active sites, making them viable drug targets, but the development of selective inhibitors has been challenging because of the non-selectivity of many deubiquitinase inhibitors and the susceptibility of the catalytic cysteine to oxidative hydrolysis (90). Unlike most deubiquitinases, COPS5 is a JAMM family zinc metallopeptidase, making it structurally distinct and potentially more amenable to selective small-molecule inhibition.

Despite the promise of COPS5 as a therapeutic target, several limitations must be addressed before clinical translation. First, CSN5i-3 is not optimized for clinical use, though our group is currently performing chemical optimization and pharmacokinetic and toxicologic profiling of CSN5i-3 derivatives. Second, NER and/or CRL repair inactivation could impair genomic integrity resulting in unintended side effects. Third, inhibition of deNEDDylation of CRLs may have context-dependent effects, potentially neutralizing anti-cancer activity by altering the balance of both tumor-suppressive and oncogenic substrates. However, it may be possible to enhance therapeutic efficacy by combining a COPS5 inhibitor with other DDR-targeting agents, such as ATR or CHK1 inhibitors, or DNA-damaging agents like PARP inhibitors. Lastly, the observed variability in COPS5 expression across patient samples underscores the need for patient stratification to identify likely responders.

Future studies should focus on refining patient selection strategies to maximize clinical impact. In summary, our study identifies COPS5 as a determinant of platinum resistance in HGSC. By modulating the NER pathway, COPS5 enables cancer cells to repair DNA damage and evade chemotherapy-induced cytotoxicity. Targeting COPS5 with CSN5i-3 is a promising therapeutic strategy that warrants further investigation to overcome platinum resistance and improve outcomes for patients with HGSC.

## METHODS

### Study Design

This study aimed to identify COPS5 as a therapeutic target to overcome platinum resistance in HGSC. Consistently, high COPS5 expression correlated with reduced DNA damage, platinum resistance, and poor clinical outcomes in ovarian cancer patients. Depleting COPS5 increases ubiquitination and degradation of essential NER and ICL repair proteins leading to impaired NER and ICL pathways. This increased carboplatin-induced DNA damage and sensitized ovarian cancer cells to carboplatin. We show that the COPS5 inhibitor CSN5i-3 synergizes with carboplatin in multiple therapy-resistant HGSC models, including patient-derived xenograft models and a syngeneic mouse model of carboplatin resistance. Together, our study shows that targeting COPS5 with CSN5i-3 is a promising therapeutic strategy that warrants further investigation to overcome platinum resistance and improve outcomes for patients with ovarian cancer.

### Sex as biological variable

Our study exclusively examined females because ovarian cancer is only relevant in females.

### Cell Lines and Culture Conditions

Sources and respective media for the 16 cell lines used in this study are detailed in Supplementary Table S5. The cell lines were maintained at 37 °C, 5% CO_2_ in a humidified incubator. The cell lines were confirmed to be negative for Mycoplasma by using the MycoAlert Mycoplasma Detection kit (LT07-318; Lonza, Rockland, ME, USA).

### siRNA Screen

The Qiagen siRNA library was used to target 98 DUBs, with four different siRNAs per gene. The Control siRNAs were: siControl (Negative Control siRNA 5 nmol, Qiagen, 1022076), siDeath (Allstars Hs Cell Death siRNA 5 nmol, Qiagen, 1027298), and siRAD51 (Silencer Pre-Designed siRNA 5 nmol, ThermoFisher, 121401). ES2 cells (3500 per well) were seeded into 96-well plates with a Multidrop, transfected in triplicate with 5 nM siRNA and the Lipofectamine RNAiMAX Transfection Reagent (ThermoFisher, 13778075) with a BiomekFX liquid handler, and incubated for 24 h. Then, cells were treated with 0.1 µM Cisplatin (Millipore Sigma, 232120) and incubated for 24 h. Cell plates were processed for DNA damage detection using the integrated screening system at the Washington University High-Throughput Screening Center. The cells were fixed with 2% paraformaldehyde in phosphate-buffered saline (PBS) for 10 minutes, washed with PBS, then permeabilized with 0.2% Triton X-100 (Thermo-Scientific, 85111) in PBS for 20 minutes. The cells were washed twice for 5 minutes, then blocked for 30 minutes with staining buffer (PBS, 0.5% BSA, 0.15% Glycine, and 0.1% Triton-X-100). Manually, cells were incubated overnight at 4 °C with γH2AX primary antibody (Millipore Sigma, 1:500) in staining buffer. Cells were washed with staining buffer for 10 minutes, then stained with secondary antibody (1:500, Alexa Fluor 488; Invitrogen) in staining buffer for 1 hour in the dark at room temperature. Cells were stained with NucBlue™ Fixed Cell ReadyProbes™ Reagent (DAPI) for 5 minutes at room temperature per the manufacturer’s instructions, then washed with staining buffer for 5 minutes. Images were collected on the InCell 2000 Analyzer high content imager and analyzed using the Multi Target Analysis Module of the InCell Analyzer 1000 Workstation Software allowing determination of cell number, quantification of γH2AX intensity and frequency (%) of γH2AX above a defined threshold. Twelve fields per well were scanned and analyzed in each plate. The negative control was siControl, no treatment, and the reference compound was siControl, 0.1 μM Cisplatin. The positive controls were as follows: siDeath, no treatment; siRAD51, 0.1 μM; siControl, 0.3 μM Cisplatin; siControl, 10 μM Cisplatin. A cut-off of three median absolute deviations (MAD) above the median was used to identify candidates that yielded greater fluorescence than the reference compound.

### Quantitative reverse transcription polymerase chain reaction

Total RNA was extracted by using the RNeasy Plus Mini Kit (Qiagen) following the manufacturer’s protocol. cDNA was synthesized with the SuperScript™ IV First-Strand Synthesis System (Thermo Fisher). Quantitative PCR was performed with Fast SYBR Green PCR Master Mix (Applied Biosystems). Gene expression was normalized to GAPDH by the ΔΔCt method. Primer sequences are provided in Supplementary Table S6.

### sgRNA cloning, lentivirus generation and transduction

COPS5 and COPS6 knockouts were generated by using a two-vector Streptococcus pyogenes (Sp) Cas9 system: pPV114-SpCas9-hygro (a gift from P. Verma) and LRG2.1 backbone (Addgene, 108098). Sequences of guides were obtained from sgRNA library [90]. sgRNAs were cloned by annealing the two complementary DNA oligos and using T4 DNA ligase to ligate them into a BsmB1-digested LRG2.1 vector. The presence of sgRNA in the plasmid DNA obtained after miniprep was verified by Sanger sequencing using the U6 primer. The sequences for primers and oligos are detailed in Supplementary Table S4. Lentiviral particles were produced by using TransIT®-LT1 Transfection Reagent (Mirus Bio) to transfect LentiX cells with lentiviral plasmids, psPAX2, and pMD2.G. Viral supernatants were collected at 48 and 72 hours post-transfection, filtered through a 0.45 μm filter, and used for infection of target cells in the presence of 8 μg/mL polybrene. Puromycin (2 µg/mL) and Hygromycin (100 µg/mL) were used for selection.

### Western blotting

Cells were lysed in urea buffer (9 M urea, 75 mM Tris-HCl, pH 7.6). Protein concentration was determined with the Bradford protein quantification assay Kit (BioRad). Equal amounts of protein (20–40 µg) were resolved by SDS-PAGE, transferred onto nitrocellulose membrane, and blocked with 10% milk or 5% BSA in tris-buffered saline with 0.1% Tween® 20. Membranes were incubated overnight at 4 °C with primary antibodies, then incubated with HRP-conjugated secondary antibodies (see Supplemental Table S7). Signal was detected via enhanced chemiluminescence (Thermo Fisher) and imaged on a ChemiDoc MP System (Bio-Rad).

### Jess Simple Western ™ Immunoblot Assay

After the indicated treatments, cells were lysed in lysis buffer (75 mM Tris-HCl, pH 7.6, 9 M urea) and clarified by centrifugation. Total protein concentration was determined with the Quick Start Bradford protein assay kit (Bio-Rad, #5000205;). The samples were analyzed on a Protein Simple Jess size-based capillary electrophoresis system (Bio-Techne, Minneapolis, MN, USA) according to the manufacturer’s instructions. Briefly, the protein lysates were diluted with sample buffer to a concentration of 1 mg/mL and then mixed with 5× fluorescent master mix at a 4:1 ratio and heated at 95 °C for 5 min. Proteins were separated by size on the 12–230 kDa Jess Separation Module (SM-W004) and detected with the Chemiluminescent Detection module. The size-separated proteins were probed with primary antibodies in antibody buffer followed by HRP-conjugated secondary antibodies (Supplementary Table S5). Data were analyzed in Compass Simple Western software, v.6.1.0 (ProteinSimple).

### Immunofluorescence

Cells were seeded at 40,000 cells per well in 8-well Nunc Lab-Tek Chamber Slide System (Thermo Fisher) and treated 24 hours later. 24 (for carboplatin, UV, or CSN5i-3) or 4 (for irradiation) hours after treatment, cells were washed with PBS, fixed with 2% paraformaldehyde for 10 minutes at 4 °C, and permeabilized with 0.2% Triton X-100 for 20 minutes at room temperature. After blocking with 0.5% BSA, 0.15% glycine, 0.1% Triton X-100 for 30 minutes, cells were incubated with primary antibodies for 1 hour at 4 °C, followed by appropriate secondary antibodies conjugated to Alexa Fluor dyes (Thermo Fisher) for 1 hour at room temperature and protected from light. Nuclei were counterstained with DAPI. Fluorescent images were captured on Leica TCS SPE inverted confocal microscope. Raw images were exported, and foci were counted with JCountPro. At least 100 cells were analyzed for each treatment group in duplicate.

### Immunohistochemistry

Tissue samples were fixed in 10% formalin, embedded in paraffin, and sectioned (5 μm). Sections were deparaffinized, rehydrated, and subjected to antigen retrieval by incubating in 10 mM citrate buffer (pH 6.0) at 95 °C for 20 minutes. Next, endogenous peroxidase was quenched in hydrogen peroxide for 15 minutes. After blocking with Biotin/Avidin (Vector Laboratories) and Protein blocking reagent (Dako), sections were incubated with primary antibodies overnight at 4 °C, and then with corresponding HRP-conjugated secondary antibodies (see Supplemental Table S5) for 1 hour at room temperature. Signals were detected with freshly made 3,3′-Diaminobenzidine (DAB) substrate solution. Sections were then counterstained with hematoxylin, dehydrated, and mounted with coverslips. Images were acquired on a Leica DM500 microscope. Immunostaining was then assessed and quantified by three independent masked researchers. Scores were assigned to samples according to the proportion and intensity of staining.

### Approval, development, and culture of primary ovarian cancer cells (POVs)

Tissues were prospectively collected for the Washington University Gynecologic Oncology biorepository (IRB #201105400 and #201706151) with informed patient consent. To establish primary cell lines, ascites was collected from patients with advanced-stage high-grade serous ovarian cancer (HGSC) and transferred to culture flasks containing 1:1 (V/V) RPMI supplemented with 20% FBS and 1% penicillin and streptomycin. After 1 to 2 weeks, attached and proliferating cells were passaged and used for experiments. Cells were discarded after 1 to 2 passages.

### Cisplatin sensitivity analysis

Data regarding COPS5 protein expression in cancer cell lines (cBioPortal database, http://www.cbioportal.org/) and corresponding cisplatin IC_50_ values were obtained from Genomics of Drug Sensitivity in Cancer [91]. GraphPad Prism 10 software was used to perform Pearson correlation analysis.

### TCGA data set analysis

Clinical data (progression-free and overall survival) and mRNA expression of COPS5 for Ovarian Serous Cystadenocarcinoma samples in TCGA PanCancer Atlas database were accessed through the public cBioPortal (http://www.cbioportal.org). The differences in survival were summarized with Kaplan–Meier curves and compared by log-rank tests in GraphPad Prism 10 and SPSS version 27.

### Patient samples/study approval

Washington University patient samples: Tissue samples were collected prospectively from patients with Stage III-IV HGSC before neoadjuvant chemotherapy as part of a national clinical trial (IRB #201407156) between December 2014 and December 2018. Chemotherapy response scores (CRS) were assigned at interval cytoreductive surgery, with CRS 3 indicating a near-pathologic complete response and CRS 1-2 classified as a poor response to platinum chemotherapy [92]. Additional samples were from tissue microarrays built in the Anatomic and Molecular Pathology Core Laboratories at Washington University (IRB# 202301067). The microarrays contained primary HGSCs collected during primary cytoreductive surgery before chemotherapy. Progression-free and overall survival were calculated from the date of surgery.

Caris Life Sciences patient samples: 12,052 high grade serous ovarian carcinoma samples were analyzed by comprehensive molecular profiling, including next-generation sequencing and whole transcriptome sequencing, performed in a CLIA/CAP/ISO15189 certified clinical laboratory (Caris Life Sciences, Phoenix, AZ, USA). In compliance with policy 45 CFR 46.101(b), this study was performed utilizing retrospective, deidentified clinical data. Therefore, this study is considered IRB exempt, and no patient consent was necessary from the patients.

Real-world overall survival information was obtained from insurance claims records, with survival durations calculated from the time of tissue collection to the last contact. Kaplan-Meier estimates were calculated for molecularly defined patient cohorts. Hazard ratios were calculated by using the Cox proportional hazards model and p values were determined with the log-rank test.

### DNA sequencing

Genomic DNA isolated from microdissected, formalin-fixed, paraffin-embedded (FFPE) samples was subjected to next-generation sequencing on the NextSeq platform (Illumina, Inc., San Diego, CA). For this, 592 whole-gene targets were enriched by using a custom-designed SureSelect XT assay (Agilent Technologies, Santa Clara, CA). All variants were detected with > 99% confidence, ascertained through allele frequency and amplicon coverage assessment. Whole-exome sequencing was performed on an Illumina Novaseq 6000 platform. For this, a hybrid pulldown of baits designed to enrich for 720 clinically relevant genes at high coverage and high read-depth was used, along with another panel designed to enrich for an additional >20k genes at a lower depth and a 500 Mb SNP backbone panel (Agilent Technologies) to help with gene amplification/deletion detection. Genetic variants identified were interpreted by board-certified molecular geneticists in alignment with American College of Medical Genetics and Genomics standards, categorizing them as ‘pathogenic’, ‘likely pathogenic’, ‘variant of unknown significance’, ‘likely benign’, or ‘benign’. When assessing mutation frequencies of individual genes, ‘pathogenic’ and ‘likely pathogenic’ were counted as mutations, and others were excluded.

### mRNA expression analysis by whole-transcriptome sequencing

RNA was extracted with the Qiagen RNA FFPE tissue extraction kit, and RNA quality and quantity were evaluated with Agilent TapeStation. Biotinylated RNA baits were hybridized to purified cDNA targets, followed by amplification of bait-target complexes in a post-capture PCR reaction. The Illumina NovaSeq 6500 was used to sequence the whole transcriptome to an average of 60 M reads. Raw data were demultiplexed by Illumina Dragen BioIT accelerator, followed by trimming, counting, removing of PCR-duplicates, and alignment to the human reference genome hg19 with the STAR aligner. Transcript counts were normalized to transcripts per million in the Salmon expression pipeline.

### Differential gene expression

Differential gene expression was analyzed by Mann-Whitney U-test for ovarian cancer patients in Caris Life Sciences.

### Colony formation assay

Cells were plated at a density of 500 cells per well (for PEO1-derived cell lines) or 300 cells per well (for OV8-derived cell lines) in six-well plates. Cells recovered for 48 or 72 hours, respectively, before treatment with carboplatin or ultraviolet (UV) irradiation. All colony formation assays presented were performed with six biological replicates (for carboplatin and UV irradiation). After treatment, cells were allowed to propagate for 11 days (for OV8-derived cell lines) or 14 days (for PEO1-derived cell lines) with media change at 6 days and 7 days, respectively. Colonies were washed with ice-cold 1x PBS and fixed in ice-cold 100% methanol for 20 minutes. They were then stained with 0.5% crystal violet solution (0.5% crystal violet, 25% methanol, and 75% deionized water) for 5 minutes. Crystal violet solution was aspirated, and plates were rinsed with DI water and placed inverted onto a paper towel and covered to dry overnight. Plates were imaged with I-bright 1500 (Invitrogen), and colonies were counted in the ColonyArea ImageJ plugin. Values were plotted as a function of drug concentration and colony area or staining intensity on GraphPad Prism.

### Cell viability assay

Cells were seeded in triplicate at 2000 cells per well in 96-well plates and treated after 24 hours as indicated in figure legends. On day 6, 20 µL of CellTiter 96 AQueous MTS Reagent (Promega, #G1112) was added to each well. After 2 to 4 hours, absorbance was measured at 490 nm on a TECAN infinite M2000PRO reader. Data was normalized to vehicle-treated controls, and background was subtracted. Values were plotted as a function of drug concentration and percent viability on GraphPad Prism, and IC_50_ values were calculated.

### Synergy analysis

Cells were treated with 8 concentrations of each drug and drug combinations in a dose–response matrix. Synergistic efficacy was assessed by using SynergyFinder 3.0 based on the Loewe drug combination model [93].

### Organoid generation and viability assays

Ascites was obtained from consenting patents with HGSC and transferred to the laboratory. Ascites (50 mL) was transferred to a 50 mL conical tube and centrifuged at 1000 rcf for 5 minutes at 4 °C. Next, cell pellets were treated with DNase (NEB; No. M0303S) and red blood cell lysis buffer (BioLegend; No. 420301). Single-cell suspensions of ascites were resuspended in 75% Cultrex (R&D Systems; No. 353300502) and 25% organoid base media [Advanced DMEM/F12 (Thermo Fisher Scientific; No. 12634028) supplemented with 1% penicillin–streptomycin (Millipore Sigma; No. P0781), 1x Glutamax (Life Technologies; No. 35050061), and 1% HEPES (Life Technologies; No. 15630080)]. The cell suspension was plated onto a 6-well plate, in approximately 40-μL droplets, and placed into a 37°C incubator for 25 minutes to solidify before adding 2 mL of organoid media [the base media, supplemented with 0.5 μM SB202190 (MedChemExpress, #HY-10295), 0.5 μM A83-01 (Sigma, #SML0788),10 ng/mL recombinant human FGF-10 (Peprotech, #100-26), 10 ng/mL recombinant human FGF-4 (Peprotech, #100-31), 100 nM β-estradiol (Sigma, #E2758), 5 mM nicotinamide (Sigma, #N0636), hEGF (Sigma-Aldrich, #E96), heregulin β-1 (Peprotech, # 100-03), hydrocortisone (Sigma-Aldrich, #H0888), and forskolin (MedChemExpress, # HY-15371). Organoid media was changed once every one to two weeks, and images were taken on day 1 after plating and periodically thereafter.

For viability assays, organoid media was pipetted up and down to dissociate organoid domes. Medium was collected into a 15 mL conical tube and centrifuged at 1000 rcf for 5 minutes at 4 °C. The supernatant was aspirated, and 1 mL of TrypLE (Thermo Fisher Scientific) was added to the pellet and vortexed briefly before incubating for 15 minutes in a 37 °C water bath. Then, the solution was centrifuged at 1000g for 5 minutes at 4 °C, and the supernatant was aspirated. The pellet was resuspended in 1 mL of organoid base media, and cells were counted on a cell counter. As described above, cells were plated at 2000 cells per 20 µL dome per well in a 48-well plate. The plate was placed in a 37 °C incubator for 25 minutes before adding 0.3 mL of organoid media into each well. Cells were allowed to propagate for 6 days before treatment. Thirteen days post treatment, CellTiter-Glo® 3D Cell Viability Assay (Promega, cat #G9681) was performed according to the manufacturer’s instructions. Luminescence was measured and graphed as a function of percent viability.

### Recovery of RNA Synthesis (RRS)

OV8 and PEO1 cells were seeded in duplicate at 40,000 cells per well in an 8-well Nunc Lab-Tek Chamber Slide System (Themo Scientific, #). Cells were treated 24 hours later with 0.1 µM CSN5i-3 or DMSO (vehicle) in media. After 24 hours, media was removed, and cells were washed with DPBS, then treated with or without 1 J/m^2^ (for PEO1) or 2 J/m^2^ (for OV8) UV irradiation. Media with vehicle or 0.1 µM CSN5i-3 was added back, and cells were left to recover for 0, 4, or 24 hours. Following recovery, the Click-iT RNA Imaging Kit (Invitrogen, #C10329) was used to measure EU incorporation. Fluorescent images were captured on a Leica DMi8 Thunder imager, and ImageJ was used for analysis.

### CometChip Assays to Measure TC-NER incision at Trabectedin-Induced Lesions

Cells were enriched at the G1 phase by incubating in growth medium supplemented with 2 μM palbociclib (hereafter referred to as the ‘working medium’) for 24 h prior to exposure to trabectedin. Following this, cells were embedded in a 30 μm COMET chip (Trevigen, 4250-096-01) and further incubated for 30 min at 37 °C in the working medium. The medium was removed, and cells were incubated in the working medium supplemented with 0.1 µM or 1µM CSN5i-3 for two hours before treating with medium with CSN5i-3 and 50 nM trabectedin for 2 h. Following trabectedin wash-out, cells were incubated in the working medium with CSN5i-3 for 0, 2 or 4 h. The CometChip was then overlaid with 6 mL of 1% low melting agarose (Trevigen, 4250-500-02), followed by cell lysis with 50 mL lysis solution (Trevigen, 4250-500-01) for overnight at 4 °C. Following lysis, DNA was unwound for 30 min twice in 250 mL alkaline solution (200 mM NaOH, 1 mM EDTA, 0.1% Triton X-100). Electrophoresis was carried out for 50 min, 1 V/cm at 4 °C in 700 mL alkaline solution. After electrophoresis, the COMET chip was neutralized for 15 min twice at 4 °C in 100 mL 0.4 M Tris pH 7.4 and equilibrated for 30 min at 4 °C in 100 mL 20 mM Tris pH 7.4, followed by staining in 50 mL 0.2 X SYBR Gold (Invitrogen, S11494) at room temperature for 2 h. The stained COMET chip was destained at room temperature in 100 mL 20 mM Tris pH 7.4 up to 1 h. Comets were imaged with 4X magnification on a fluorescence microscope (Olympus, BX53). % DNA in tail was quantified with Comet analysis software (Trevigen, 4260-000-CS) [51].

### DNA Fiber Assay

OVCAR8 cells were seeded in 10-cm dishes. After settling overnight, cells were treated with 0.1 µM CSN5i-3 or vehicle for 24 hours, washed with DPBS and treated with the same treatment media, with the addition of 50 µM 5-Chloro-2’-deoxyuridine (CldU, Millipore Sigma), for 20 minutes. Cells were washed twice with DPBS and treated with the above treatments, with the addition of 500 µM carboplatin and 5-Iodo-2’-deoxyuridine (IdU, Millipore Sigma), for 60 minutes, then washed twice again. Cells were detached from plates with 0.05% Trypsin EDTA, pelleted, and resuspended in ice-cold PBS. Cells were diluted to a concentration of 500,000 cells/mL in ice-cold PBS with 0.1% BSA. 4 µL of each sample was placed on charged glass slides (Globe Scientific) and incubated at room temperature for 4 minutes. 6 µL lysis buffer (200 mM Tris-HCl pH 7.5, 50 mM EDTA, 0.5% SDS in ddH_2_O) was added to each sample and pipetted 5 times to mix before incubating at room temperature for 4 minutes. Slides were then placed at a 20° angle to allow lysed samples to migrate to the ends. Slides were allowed to dry and fixed in 3:1 methanol-acetic acid for five minutes. To stain the DNA fibers, slides were washed twice with PBS for 5 minutes each and incubated in 2.5N HCl for 1 hour at room temperature. Slides were washed 3 times with PBS and incubated in blocking buffer (2% BSA, 0.2% Tween-20 in PBS) for 1 hour. Slides were then incubated in primary antibodies (1:200, rat anti-BrdU, Abcam; 1:200, mouse anti-BrdU, BD Biosciences) at room temperature for 2 hours. Slides were washed with PBST (0.2% Tween-20 in PBS) 4 times and with PBS once, then incubated in secondary antibodies (1:200, anti-rat Alexa Fluor 488, Invitrogen; 1:200, anti-mouse Alexa Fluor 568, Invitrogen) at room temperature for 1 hour. Slides were washed 4 times with PBST, and cover glasses were affixed with mounting medium (Invitrogen, ProLong Gold). Fluorescent images were captured on a Leica TCS SPE inverted confocal microscope, and fiber lengths were measured using ImageJ and used to calculate IdU/CldU ratio. All analysis was blinded and a minimum of 100 fibers were measured for each condition.

### TUBEs-Mass Spectrometry for Identification and Analysis of the Ubiquitin-Proteome

Identification of ubiquitylated proteins was performed using TUBEs-LC-MS/MS technology by LifeSensors Inc (Malvern, PA). OV8 cells were seeded at 1,000,000 cells per well and incubated overnight. Cells were treated with 0.5 µM CSN5i-3, DMSO (vehicle), or combination of CSN5i-3 and MG132 for 8 hrs. Briefly, cell lysates were incubated with tandem ubiquitin-binding entities (TUBEs), and enriched polyubiquinated fractions were eluted and lyophilized. Lyophilized elutes from TUBE pulldowns were resuspended, and the entire sample was run 0.5 cm into a preparative SDS-gel. The excised gel regions were reduced with TCEP, alkylated with iodoacetamide, and digested with trypsin. Tryptic digests were analyzed using a 40-min LC-MS/MS OT-AT data-dependent acquisition on the Thermo Orbitrap Astral mass spectrometer. MS/MS data were analyzed using Thermo Proteome Discoverer v3.2.0.450. Spectra were searched against the SwissProt human database (UP000005640, downloaded 05-04-2025). Gene Set Enrichment Analysis (GSEA) was performed to identify significantly enriched Gene Ontology: Biological Process (GO:BP) terms using the MSigDB v2025.1. Hs human gene set collection, specifically the C5: GO Gene Sets and the GO:BP sub-collection. Gene sets were retrieved from the Molecular Signatures Database (MSigDB), and the enrichment analysis was conducted using the official GSEA platform. A ranked list of genes was generated based on log2 fold change, and the top 500 genes were selected as input for the enrichment analysis. From the results, the top 100 significantly enriched GO:BP terms were selected based on adjusted p-values (FDR q-values). Of these, the top 30 pathways ranked by gene ratio were visualized using bubble plot to highlight the key biological processes involved.

### Animal Study

All mouse experiments were approved by the Institutional Animal Care and Use Committee in accordance with the Animal Welfare Act, the Guide for the Care and Use of Laboratory Animals and NIH guidelines (IACUC protocol No. 24-0238) at Washington University in St. Louis. All procedures were performed in accordance with the guidelines of the American Association for Accreditation for Laboratory Animal Care and the US Public Health Service Policy on Human Care and Use of Laboratory Animals. Mice (6 to 7 weeks old, female) NU/J (strain 002019) or C57BL/6J (strain 000664) mice were purchased from Jackson Laboratories. Mice were weighed twice per week and examined daily for signs of distress.

NU/J mice were intraperitoneally inoculated with 4×10^6 sg*Neg*, sg*COPS5*, or sg*COPS6* OVCAR8 cells. Treatment was initiated fourteen days after inoculation. At this point, animals were randomized into two treatment groups: vehicle (n=10 per group), and 60mg/kg carboplatin (SelleckChem, Cat# S1215, n=10 per group). Carboplatin (dissolved in Dulbecco’s phosphate buffered saline with MgCl_2_ and CaCl_2_) and vehicle (Dulbecco’s phosphate buffered saline with MgCl_2_ and CaCl_2_) were administered intraperitoneally once a week for two weeks. Following treatment period, mice were euthanized, and tumor burden was assessed. Tumor samples were collected, fixed in formalin, embedded in paraffin wax, and sectioned in 4 µm thick slices for immunofluorescent analysis.

C57BL/6J mice were intraperitoneally inoculated with 1×10^7 ID8-CPR cells. Treatment was initiated twenty-one days after inoculation. At this point, animals were randomized into four treatment groups: (1) Vehicle 1 and Vehicle 2 (n=15), (2) Vehicle 1 + 20 mg/kg Carboplatin (SelleckChem, Cat# S1215, n=15), (3) Vehicle 2 + 50 mg/kg CSN5i-3 (MedChemExpress, Cat# HY-112134, n=15), and (4) 20 mg/kg Carboplatin + 50mg/kg CSN5i-3 (n=15). CSN5i-3 was formulated in Vehicle 1 – 100 mM citrate buffer, pH 3, PEG300, Kolliphor HS15, and 1 N HCl (56:30:10:4%w/w). Carboplatin was dissolved in Vehicle 2 - Dulbecco’s phosphate buffered saline with MgCl_2_ and CaCl_2_. Carboplatin and Vehicle 2 were administered intraperitoneally once per week for two weeks. CSN5i-3 and Vehicle 1 were administered via oral gavage five days per week for two weeks. Following treatment period, 4-6 mice in each treatment group were monitored for survival, and the rest of mice were euthanized, and tumor burden was assessed. Tumor samples were collected, fixed in formalin, embedded in paraffin wax, and sectioned in 5 µm thick slices for immunohistochemistry.

### Statistical analysis

Continuous variables were compared by non-parametric tests, including Wilcoxon/Mann-Whitney U tests, and parametric tests, such as independent Student’s t-tests and one-way ANOVA with Tukey’s multiple comparison tests when appropriate. Pearson correlation analysis was used to assess relationships between variables. Multiple comparisons were adjusted by using the Benjamini–Hochberg method, with an adjusted p-value (q-value) < 0.05 considered significant. Traditional statistical analyses were conducted in GraphPad Prism 9 and SPSS version 27. All experiments were performed at least twice, each in triplicate, unless otherwise noted. Statistical significance was set at P < 0.05, and two-tailed 95% confidence intervals are reported.

## List of Supplementary Materials

Materials and Methods

Fig S1 to S7 for multiple supplementary figures

Tables S1 to S7 for multiple supplementary tables

## Acknowledgements

We wish to acknowledge all the patients who have generously donated their cancer specimens to make this work feasible. We would also like to acknowledge Deborah Frank, PhD for her manuscript editing, the High-Throughput Screening Core at Washington University for their services, and Mr. Glen Huns for their support of this project.

## Funding

Damon Runyon Cancer Research Foundation

The Reproductive Scientist Development Program (RSDP) supported by the Gynecologic Oncology Group Foundation

The NCI Early-Stage Surgeon Scientist Program

The Victoria’s Secret Global Fund for Women’s Cancers Career Development Award in partnership with Pelotonia and the American Association for Cancer Research (AACR)

The Doris Duke COVID-19 Fund to Retain Clinical Scientists (CFRCS)

The Foundation for Women’s Cancer.

## Author Contributions

Conceptualization: EL, MMM

Methodology: EL, ML, KR, MMM

Investigation: EL, ML, KR, LNVB, MM, JB, MB, RD, CS, EG, PKT

Visualization: EL, ML, KR, MMM

Funding acquisition: MMM

Project administration: MMM

Supervision: EL, MMM

Writing – original draft: EL, ML, KR, MMM

Writing – review & editing: EL, ML, KR, LNVB, JB, AS, MM, JB, MB, JP, RD, CS, AO, EG, PKT, NS, SW, MJO, BS, LK, CM, ARH, PT, DM, MP, ISH, MXGI, KF, PV, ODS, NM, DK, MMM

## Data availability

The datasets generated and/or analyzed during the current study are available from the corresponding author on reasonable request. The de-identified sequencing data from Caris Life Sciences cannot be publicly shared because of patient privacy. Qualified researchers can apply for access by contacting Caris Life Sciences and signing a data usage agreement. Other questions regarding the data of this study should be directed to the corresponding author, M.M.

