## Supplementary Table S2 for "Targeting the COP9 signalosome overcomes platinum resistance in ovarian cancer through two distinct genome stability mechanisms"

Sequence identity between the *H. sapiens* and *M. musculus* orthologues CSN subunits.

| *H. Sapiens* | GPS1 | COPS2 | COPS3 | COPS4 | COPS5 | COPS6 | COPS7a | COPS7b | COPS8 | COPS9 |
| --- | --- | --- | --- | --- | --- | --- | --- | --- | --- | --- |
| vs. | vs. | vs. | vs. | vs. | vs. | vs. | vs. | vs. | vs. | vs. |
| *M. Musculus* | Gps1 | Cops2 | Cops3 | Cops4 | Cops5 | Cops6 | Cops7a | Cops7b | Cops8 | Cops9 |
| Sequence identity (%) | 97.1 | 100 | 99.5 | 99.5 | 99.4 | 98.1 | 98.9 | 98.1 | 95.2 | 98.2 |
