## Supplemental Figures for "Targeting the COP9 signalosome overcomes platinum resistance in ovarian cancer through two distinct genome stability mechanisms"

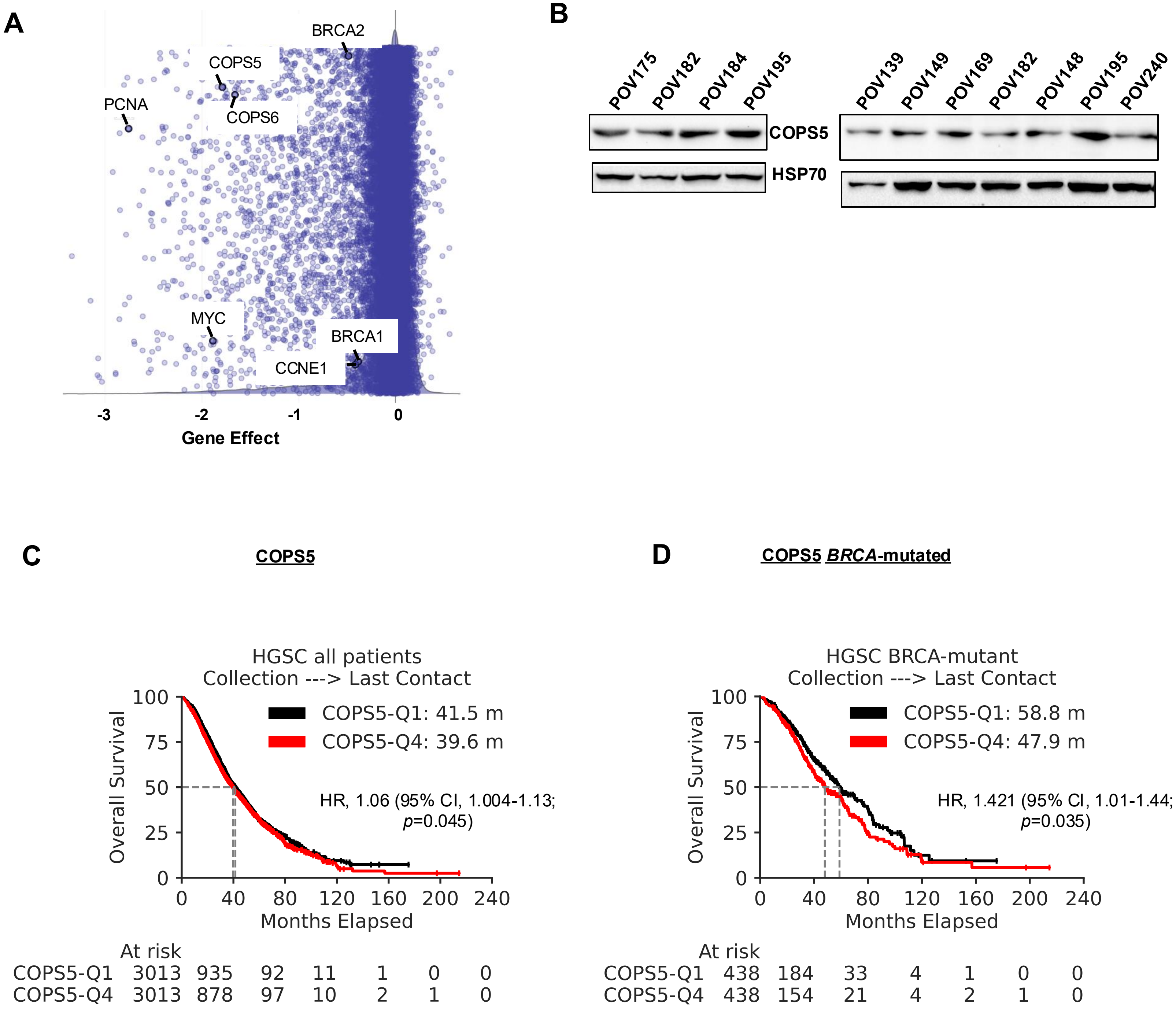


**Fig. S1. Evaluation of COPS5 as a critical gene in ovarian cancer cells and patient survival** (**A**) Cancer dependency scores of common target genes derived from CRISPR knockout screen datasets (DepMap) across various ovarian cancer cell lines. Negative scores indicate cell growth inhibition upon gene knockout. (**B**) COPS5 protein expression in indicated POVs, as determined by Western blotting, with HSP70 expression serving as the loading control and analyzed by densitometry. (**C**) Kaplan-Meier survival analysis comparing the OS between ovarian cancer patients with high (upper quartile) and low (lower quartile) *COPS5* mRNA expression Median survival (months, mean ± SD), HR with 95% CIs and P value for log-rank test are shown. (**D**) Kaplan-Meier survival analysis comparing the OS between BRCA-mutated ovarian cancer patients with high (upper quartile) and low (lower quartile) *COPS5* mRNA expression Median survival (months, mean ± SD), HR with 95% CIs and P value for log-rank test are shown.

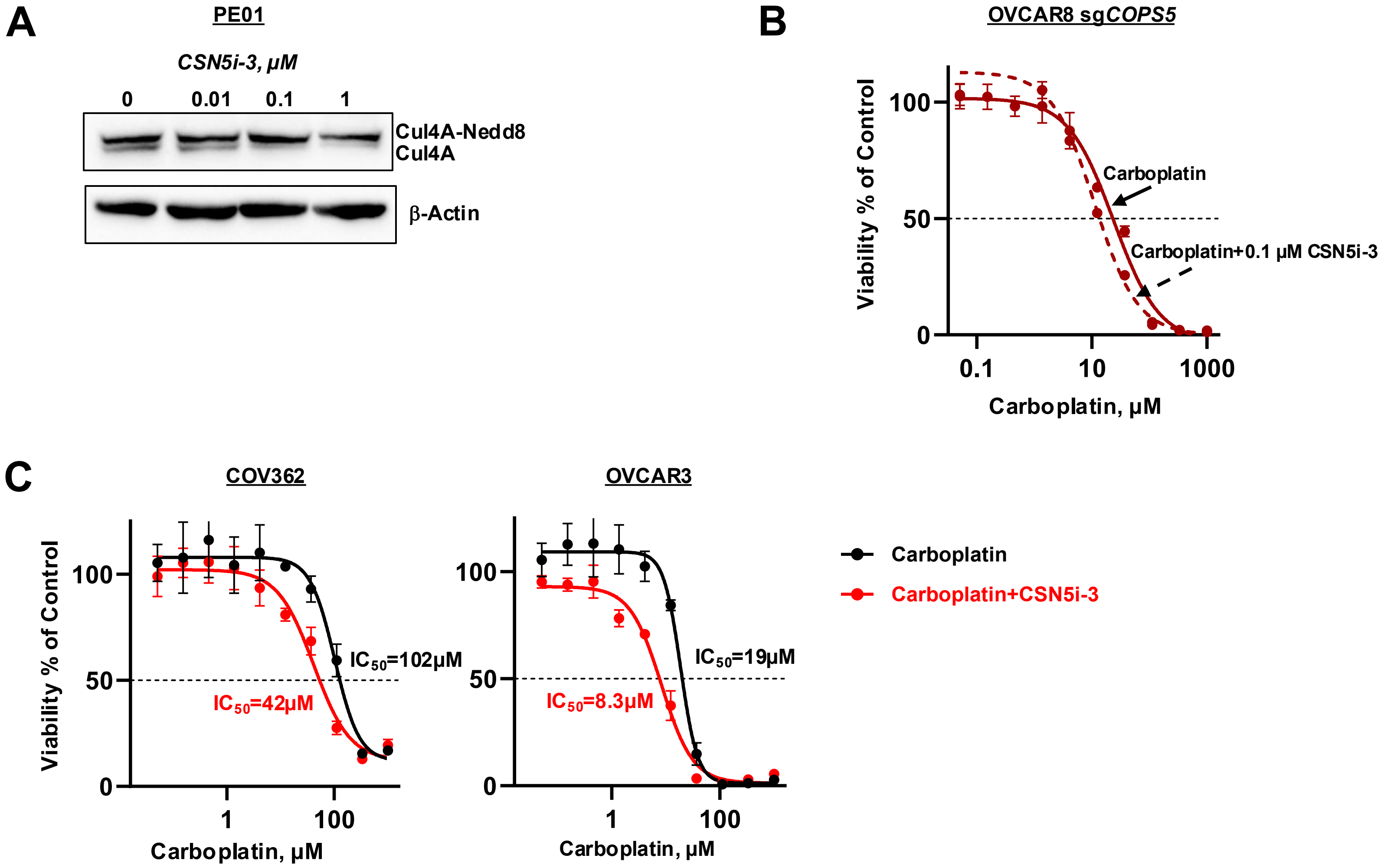


**Fig. S2. CSN5i-3 modulates Cullin4A expression and increases carboplatin sensitivity in ovarian cancer cells** (**A**) Western blots showing Cullin4A expression after treatment of PE01 cells with indicated concentrations of CSN5i-3, with β-actin expression serving as the loading control. (**B-C**) Viability of indicated cells treated with increasing doses of carboplatin alone or in combination with 0.1 μM CSN5i-3, as measured by MTS assay. Dose–response curves represent normalized viability (means ± SD, n = 3) fitted in log(inhibitor) vs. response Hill variable slope model. $IC_{50}$ values (means ± SD, n = 3) are indicated.

**A**

```
Q92905|CSN5-H.Sapiens     1 MAASGSGMAQKTWELANNMQEAQSIDEIYKYDKKQQQEILAAKPWTKDHHYFKYCKISAL
O35864|CSN5-M.Musculus    1 MAASGSGMAQKTWELANNMQEAQSIDEIYKYDKKQQQEILAAKPWTKDHHYFKYCKISAL

Q92905|CSN5-H.Sapiens    61 ALLKMVMHARSGGNLEVMGLMLGKVDGETMIIMDSFALPVEGTETRVNAQAAAYEYMAAY
O35864|CSN5-M.Musculus   61 ALLKMVMHARSGGNLEVMGLMLGKVDGETMIIMDSFALPVEGTETRVNAQAAAYEYMAAY

Q92905|CSN5-H.Sapiens   121 IENAKQVGRLENAIGWYHSHPGYGCWLSGIDVSTQMLNQQFQEPFVAVVIDPTRTISAGK
O35864|CSN5-M.Musculus  121 IENAKQVGRLENAIGWYHSHPGYGCWLSGIDVSTQMLNQQFQEPFVAVVIDPTRTISAGK

Q92905|CSN5-H.Sapiens   181 VNLGAFRTYPKGYKPPDEGPSEYQTIPLNKIEDFGVHCKQYYALEVSYFKSSLDRKLLEL
O35864|CSN5-M.Musculus  181 VNLGAFRTYPKGYKPPDEGPSEYQTIPLNKIEDFGVHCKQYYALEVSYFKSSLDRKLLEL

Q92905|CSN5-H.Sapiens   241 LWNKYWVNTLSSSSLLTNADYTTGQVFDLSEKLEQSEAQLGRGSFMLGLETHDRKSEDKL
O35864|CSN5-M.Musculus  241 LWNKYWVNTLSSSSLLTNADYTTGQVFDLSEKLEQSEAQLGRGSFMLGLETHDRKSEDKL

Q92905|CSN5-H.Sapiens   301 AKATRDSCKTTIEAIHGLMSQVIKDKLFNQINIS
O35864|CSN5-M.Musculus  301 AKATRDSCKTTIEAIHGLMSQVIKDKLFNQINVA
```

**B**

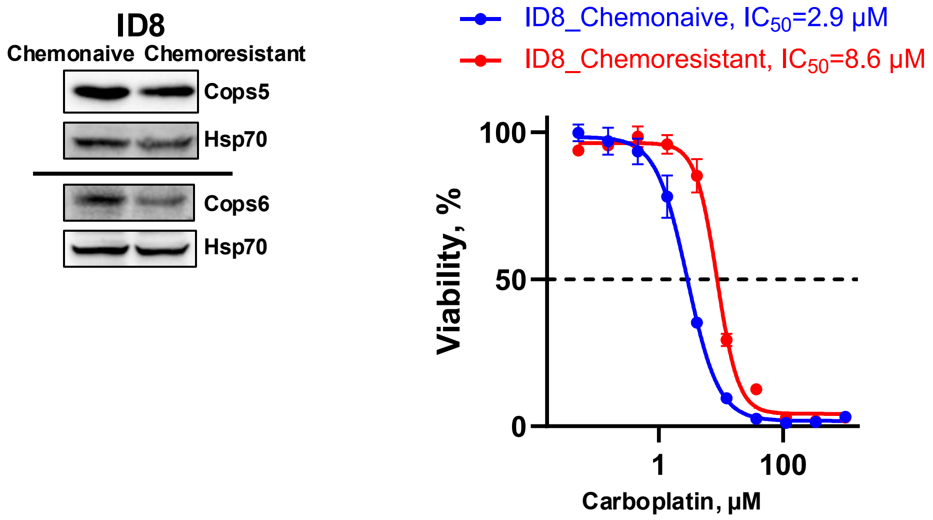


**Fig. S3. COPS5 domain conservation across species and characterization of chemoresistant ID8-CPR ovarian cancer cells** (**A**) Amino acid sequence alignment, produced by BLAST, of *H. sapiens* and *M. musculus* COPS5 proteins. The one-letter amino acid code in bold and the outlined letters represent Mpr1-Pad1-N-terminal (MPN) domain (aa 55-192) and Jab1 MPN domain metalloenzyme (JAMM) motif (aa 138-151), respectively, at 100% sequence similarity. (**B**) COPS5 and COPS6 protein expression in indicated cells, as determined by Western blotting, with HSP70 expression serving as the loading control. Viability of indicated cells treated with increasing doses of carboplatin, as measured by MTS assay. Dose–response curves represent normalized viability (means ± SD, n = 3) fitted in log(inhibitor) vs. response Hill variable slope model. $IC_{50}$ values (means ± SD, n = 3) are indicated.

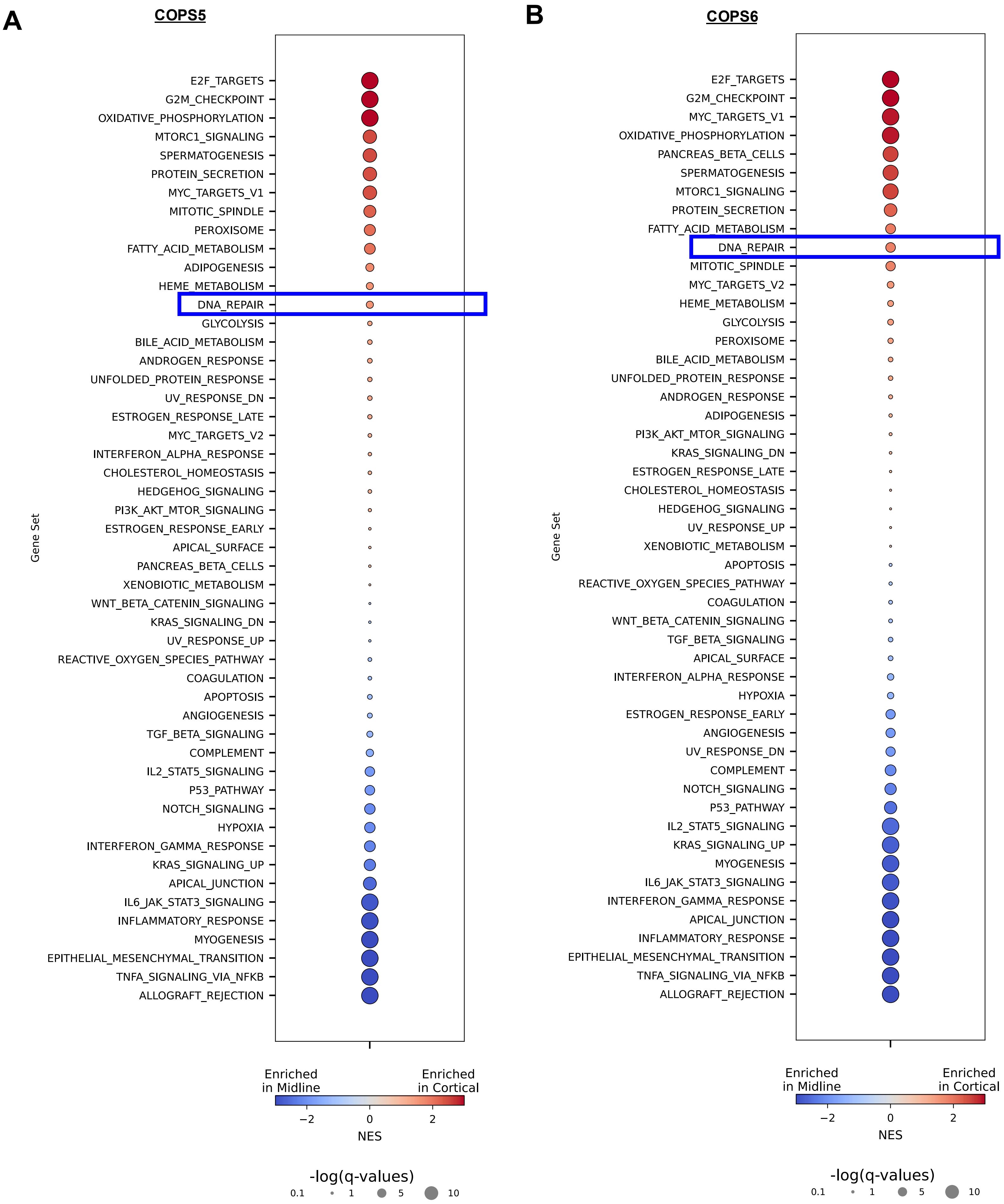


**Fig. S4. Transcriptomic data from ovarian cancer samples.** Pathway enrichment analysis. Top-ranked pathways associated with differentially expressed genes between (**A**) COPS5-High (upper quartile) versus COPS5-Low (lower quartile) and (**B**) COPS6-high versus COPS6-low in the Caris dataset.

A

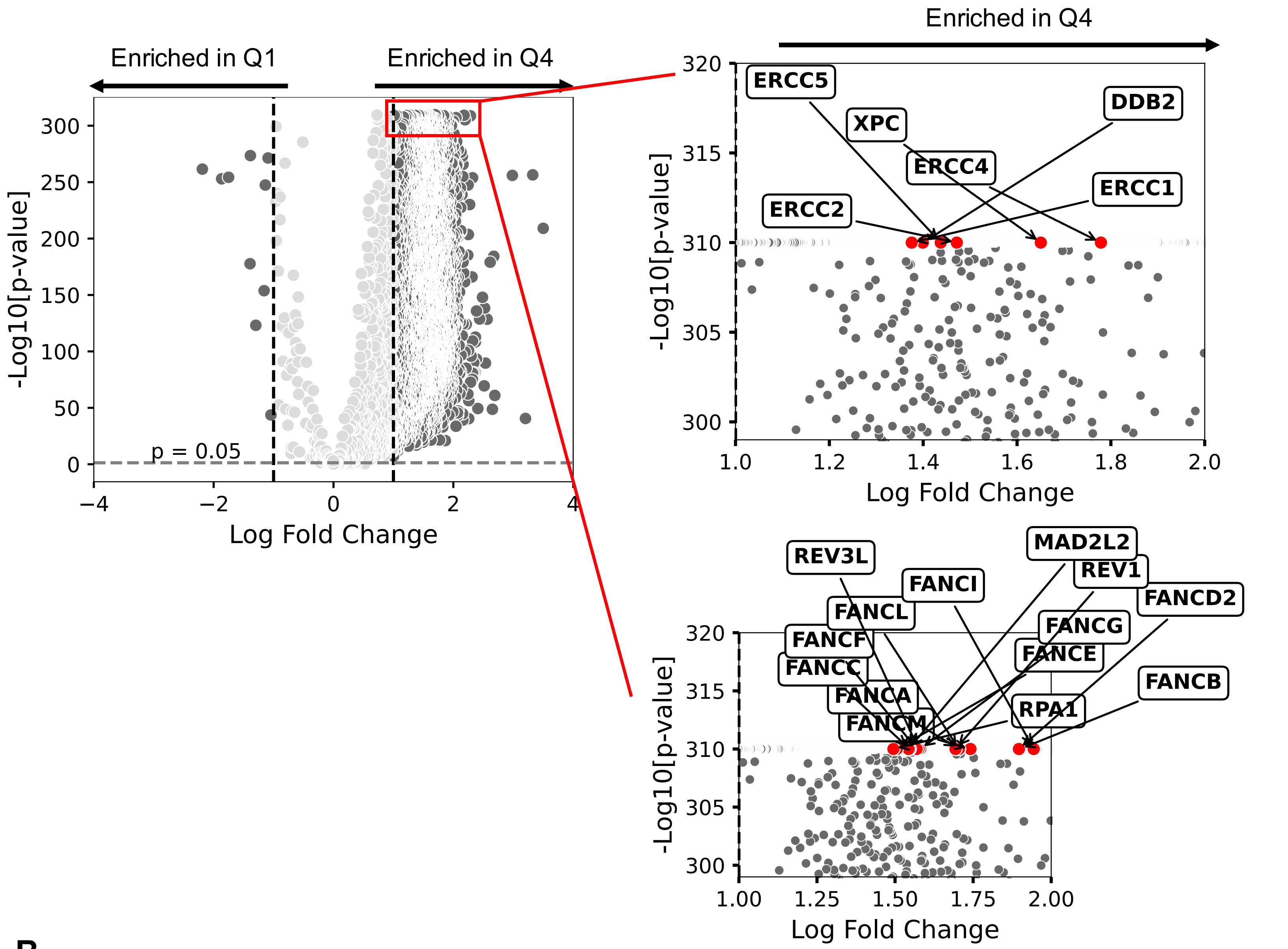

Enriched in Q1
Enriched in Q4
-Log10[p-value]
Log Fold Change
p = 0.05
ERCC5
XPC
ERCC4
DDB2
ERCC1
ERCC2
REV3L
FANCI
MAD2L2
REV1
FANCD2
FANCL
FANCF
FANCC
FANCA
FANCM
FANCG
FANCE
RPA1
FANCB


B

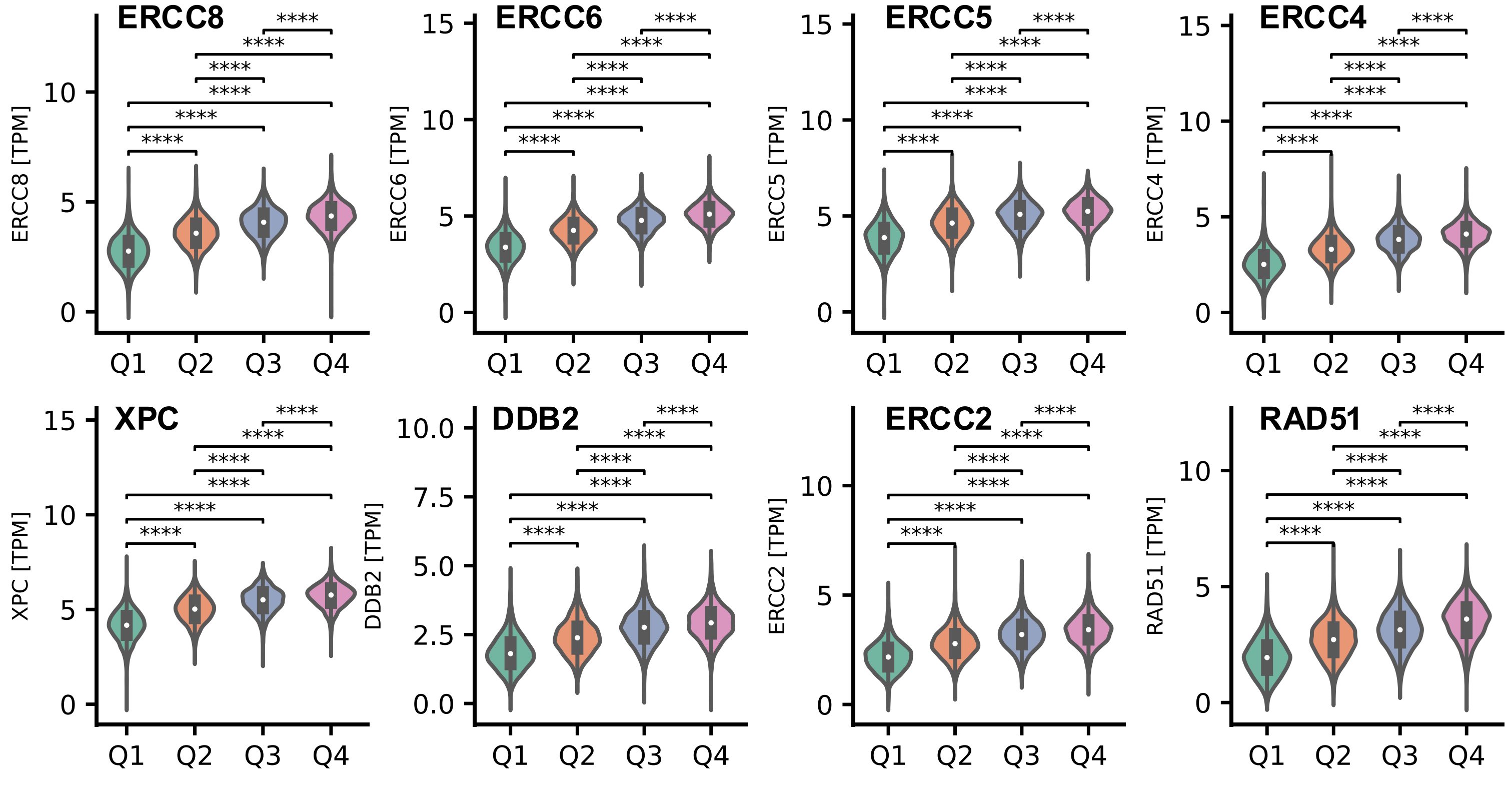

ERCC8
ERCC6
ERCC5
ERCC4
XPC
DDB2
ERCC2
RAD51
ERCC8 [TPM]
ERCC6 [TPM]
ERCC5 [TPM]
ERCC4 [TPM]
XPC [TPM]
DDB2 [TPM]
ERCC2 [TPM]
RAD51 [TPM]
Q1
Q2
Q3
Q4
****

**B**

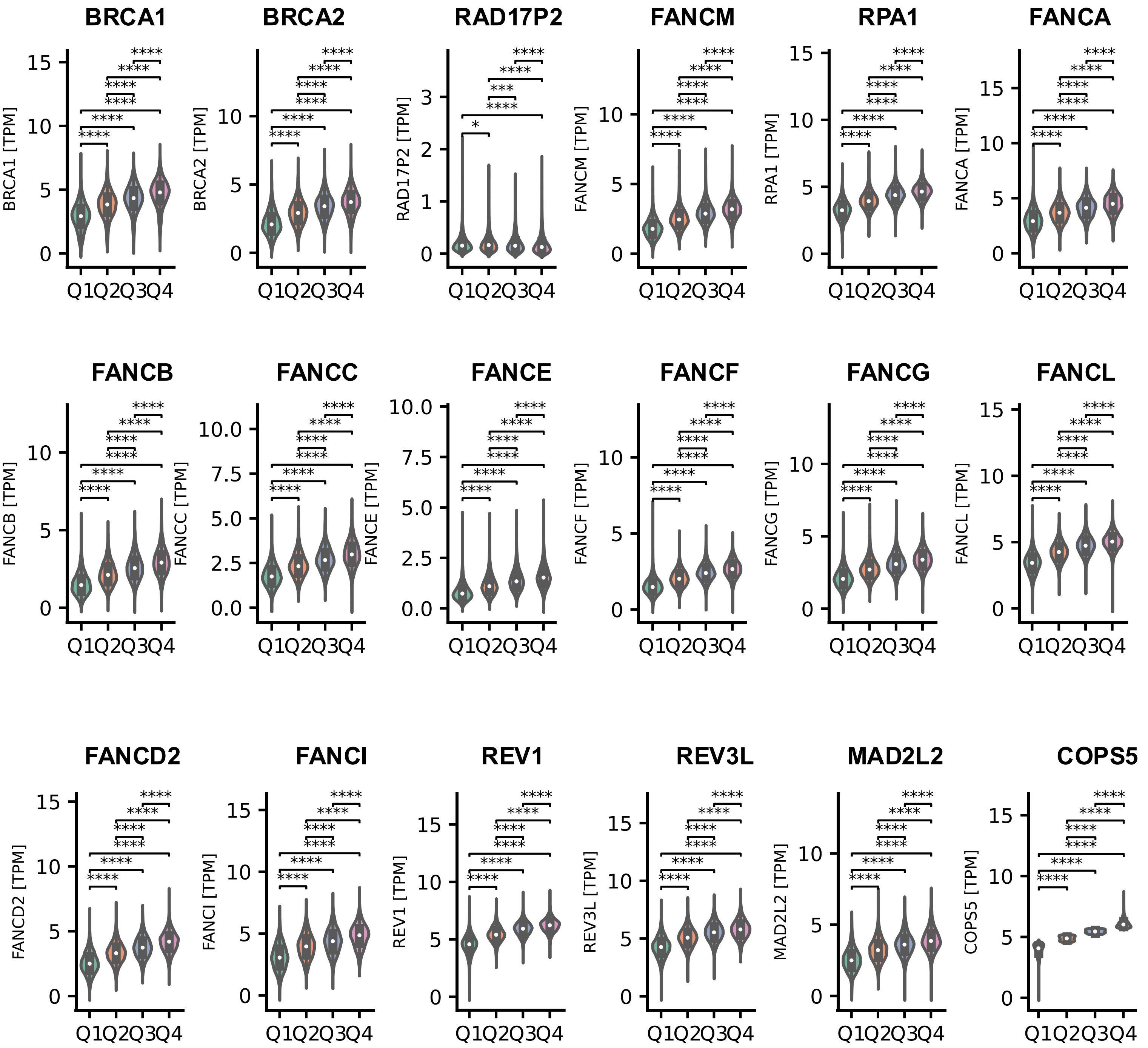


**Fig. S5. Transcriptomic data indicating enrichment of DNA repair protein expression in COPS5-high tumors** (**A**) Volcano and enhanced volcano plots (inserts) of differentially expressed genes in COPS5-High versus COPS5-Low tumors in the Caris dataset. Results are expressed as fold change versus P value. (**B**) Violin plots showing mRNA levels of indicated genes based on COPS5 expression from low (quartile 1) to high (quartile 4) in the Caris dataset. Mann-Whitney U test is used in A and B.

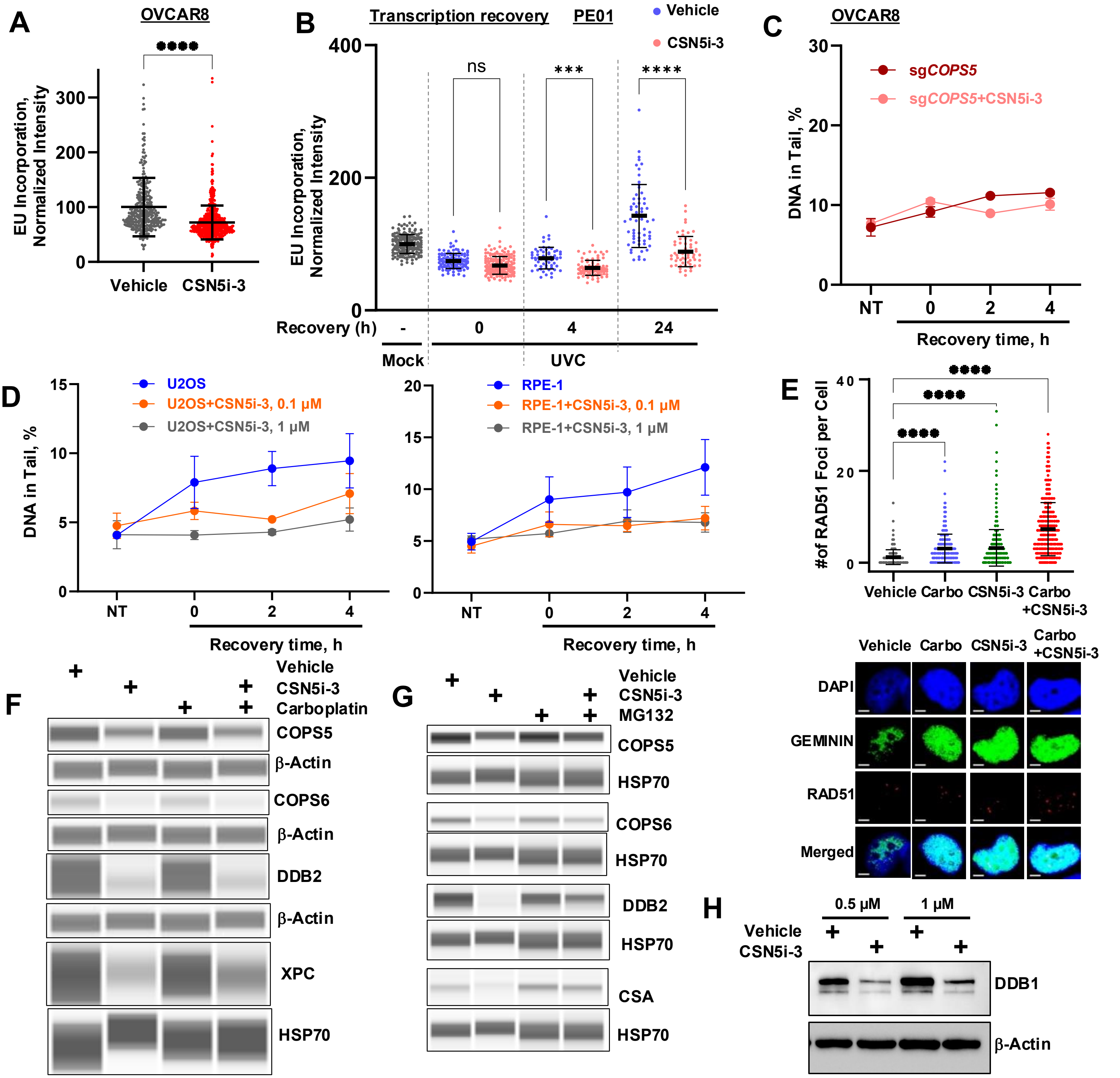


**Fig. S6. Effect of CSN5i-3 on transcription recovery, RAD51 foci formation, and protein expression in ovarian cancer cells** (**A**) Scatter plot showing nascent transcripts labelled with EU and assessed at 24 hours after vehicle or 0.1 μM CSN5i-3 administration in OVCAR8 cells. (**B**)Transcription recovery following UVC-induced DNA damage in CSN5i-3-treated PEO1 cells. (**C**) Summary and statistical analysis of ssDNA breaks analyzed by alkaline COMET chip assay in sgCOPS5 and sgCOPS5 + CSN5i-3 in OVCAR8 cells treated with CSN5i-3 (2h) and then with trabectedin (50 nM, 2 h) and allowed to recover for up to 4 h. Mean ± SEM of three biological replicates. (**D**) Summary and statistical analysis of ssDNA breaks analyzed by alkaline COMET chip assay in U2OS and RPE-1 cells treated with CSN5i-3 (2h) and then with trabectedin (50 nM, 2 h) and allowed to recover for up to 4 h. Mean ± SEM of three biological replicates. (**E**) Quantification and representative images of RAD51 foci in OVCAR8 treated with vehicle, carboplatin, CSN5i-3, and carboplatin-CSN5i-3 combination. Scale bars, 5 μm. n = 3 replicates, ****, $p < 0.0001$ based on multiple two-sided Mann–Whitney tests. (**F**) Jess Simple Western blots showing expression of indicated proteins in OVCAR8 cells treated with vehicle, CSN5i-3, or carboplatin with β-actin or HSP70 as the loading control.(**G**) Jess Simple Western blots showing expression of indicated proteins in OVCAR8 cells treated with vehicle, CSN5i-3, MG132, or CSN5i-3-MG132, with HSP70 as the loading control. (**H**) Western blot showing expression of DDB1 in OVCAR8 cells treated with vehicle or CSN5i-3 at indicated concentrations, with β-actin as the loading control.

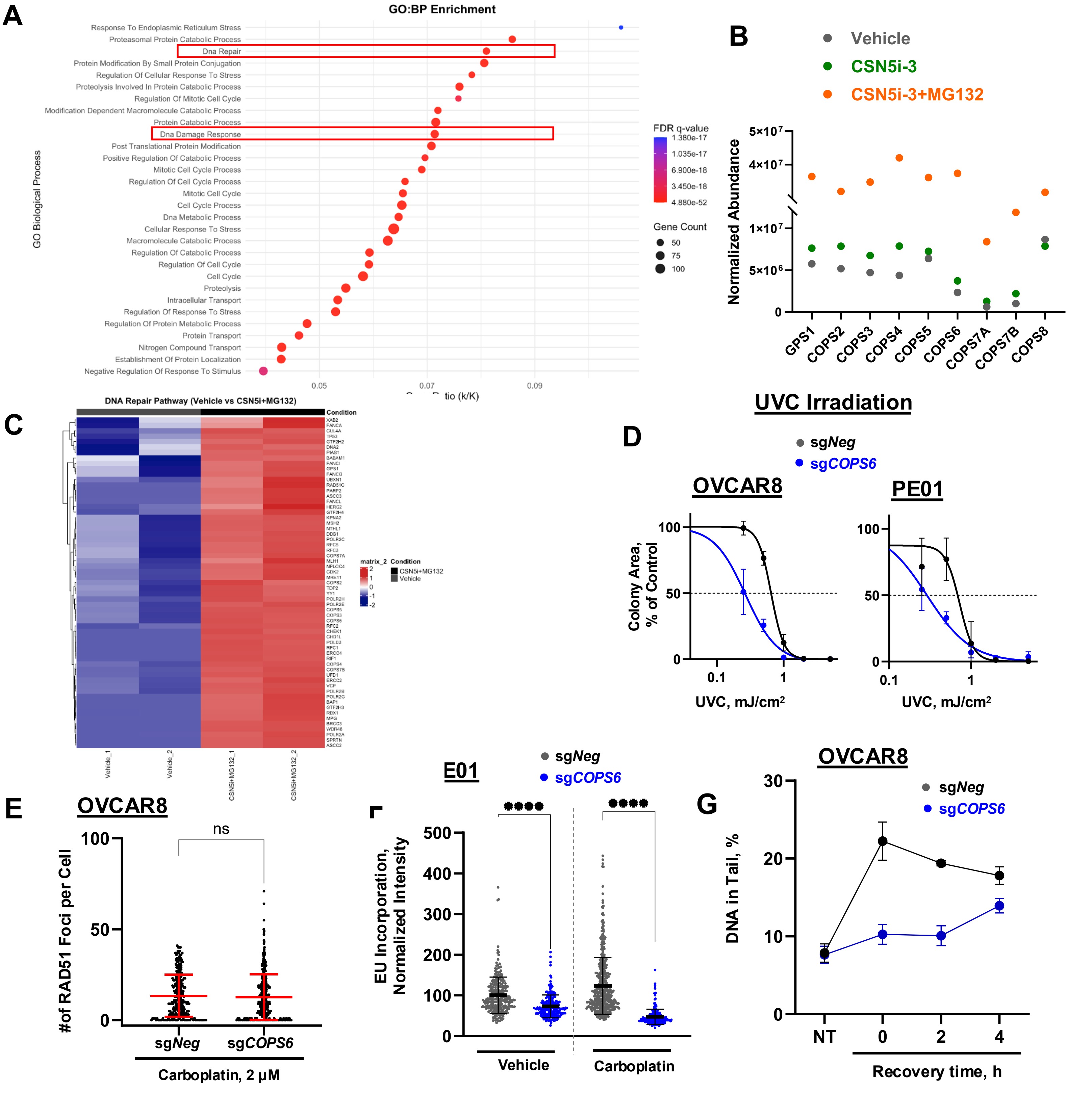


**Fig. S7. Loss of COPS6 phenocopied the effects of COPS5 loss** (**A**) GSEA on differentially abundant ubiquitinated proteins using MsigDB gene ontology: biological process gene sets, showing enrichment of DNA repair pathways. Pathways are ranked by adjusted P values <0.05. (**B**) Abundance of COP9 subunits by TUBE-based mass spectrometry ubiquitin proteomics. (**C**) Heatmap of normalized protein abundance for DNA repair pathway. (**D**) Viability of indicated cells treated with increasing doses of UVC alone as measured by clonogenic assay. Dose–response curves represent normalized viability (mean ± SD, n = 3) fitted in log(inhibitor) vs. response Hill variable slope model. (**E**) Quantification of RAD51 foci in OVCAR8 sg*Neg* and sg*COPS6* cells treated with carboplatin. (**F**) Scatter plot showing nascent transcripts labelled with EU and assessed after 24 hours after vehicle or carboplatin administration in PE01 sg*Neg* and sg*COPS6* cells. Dots indicate normalized nuclear EU intensity per cell. Data are mean ± SD from at least 110 cells. (**G**) Summary and statistical analysis of ssDNA breaks analyzed by alkaline COMET chip assay in OVCAR8 sg*Neg* and sg*COPS6* cells treated with trabectedin (50 nM, 2 h) and allowed to recover for up to 4 h. Mean ± SEM of three biological replicates. Each dot represents DNA in tail (%) of a comet analyzed. Mean ± SEM of three biological replicates. Each dot represents DNA in tail (%) of a comet analyzed.
